# Linker Length and Composition within Disordered Binding Motifs modulates the Avidity and Reversibility of a Multivalent Protein Interaction Switch

**DOI:** 10.1101/2025.06.06.658374

**Authors:** Kiran Sankar Chatterjee, Maria A. Martinez-Yamout, H. Jane Dyson, Peter E. Wright

**Affiliations:** Department of Integrative Structural and Computational Biology and Skaggs Institute of Chemical Biology, Scripps Research Institute, 10550 North Torrey Pines Road, La Jolla, California 92037

**Author notes:** Author for correspondence: Peter E. Wright.

## Abstract

Intrinsically disordered proteins that mediate the cellular transcriptional response to hypoxia play important roles in turning on and turning off oxygen stress genes. In particular, the feedback inhibitor CITED2 operates a unidirectional switch that efficiently terminates the hypoxic response by displacing the C-terminal activation domain of the hypoxia-inducible factor HIF-1α from its complex with the TAZ1 domain of the transcriptional coactivators CBP and p300. Unidirectionality of the switch arises from subtle allosteric conformational changes in TAZ1 and from differences in the strength of thermodynamic coupling between the TAZ1-binding motifs in the multivalent HIF-1α and CITED2 activation domains. To investigate the role of binding cooperativity, we mutated a linker sequence in the HIF-1α activation domain to enhance or reduce the thermodynamic coupling between its TAZ1-binding motifs. An efficient native-gel assay shows that certain linker mutations enhance the affinity of HIF-1α for TAZ1, and fluorescence anisotropy competition and NMR measurements show that these mutants can compete with CITED2 for TAZ1 more effectively than wild-type HIF-1α. The wide range of mutants, which include insertion, deletion, replacement and scrambling of residues in the linker, provide insights into the molecular basis for the exquisite tuning of the hypoxic switch: the TAZ1 affinity and consequent CITED2 competition enhancement depends both on the flexibility of the linker sequence (particularly the presence of glycine residues) and the unfavorable electrostatic interactions of a highly conserved arginine side chain in the center of the linker with an electropositive surface of TAZ1.

## Introduction

Multivalent interactions mediated by intrinsically disordered proteins (IDPs) play a major role in cellular signaling networks.^*1–3*^ Multivalency and conformational dynamics are among the key properties of IDPs that determine binding affinity/avidity, mediate cross talk between regulatory pathways, facilitate competition for binding to common sites on target proteins, and enable allosteric regulation of signaling networks.^*4,5*^

The cellular response to hypoxia in vertebrates provides a paradigmatic example of the function of multivalent IDPs in controlling key regulatory processes. In normoxic cells, the hypoxia inducible transcription factor HIF-1α is hydroxylated and targeted for degradation.^*6*^ Under conditions of hypoxia, hydroxylation is suppressed and HIF-1α activates transcription of oxygen stress genes by recruiting the transcriptional coactivators CBP/p300 through interactions with their TAZ1 domain.^*7,8*^ CITED2, which is expressed under the control of HIF-1α, functions as a negative feedback regulator that down-regulates the hypoxic response by competing with HIF-1α for binding to the CBP/p300 TAZ1 domain.^*9,10*^ Both HIF-1α and CITED2 bind TAZ1 through their intrinsically disordered C-terminal activation domains (CTAD). CITED2 displaces HIF-1α from its complex with TAZ1 by a facilitated dissociation process, forming a hypersensitive allosteric switch that attenuates the hypoxic response.^*10*^ The switch operates through the efficient displacement of HIF-1α from its complex with TAZ1 by CITED2, while HIF-1α is ineffective in rebinding to displace CITED2, ensuring that the switch is unidirectional.

Multivalency of the disordered HIF-1α and CITED2 activation domains and differences in their conformational dynamics when bound to TAZ1 are central to the functioning of the unidirectional switch. Both activation domains fold to form helical structure upon binding to TAZ1.^*11–14*^ The HIF-1α activation domain binds TAZ1 via an LPQL interaction motif and two helices, termed αB and αC, which are connected by a flexible linker (Figure 1).^*11,12,15*^ An additional helix (αA) interacts transiently with TAZ1.

**Figure 1.**
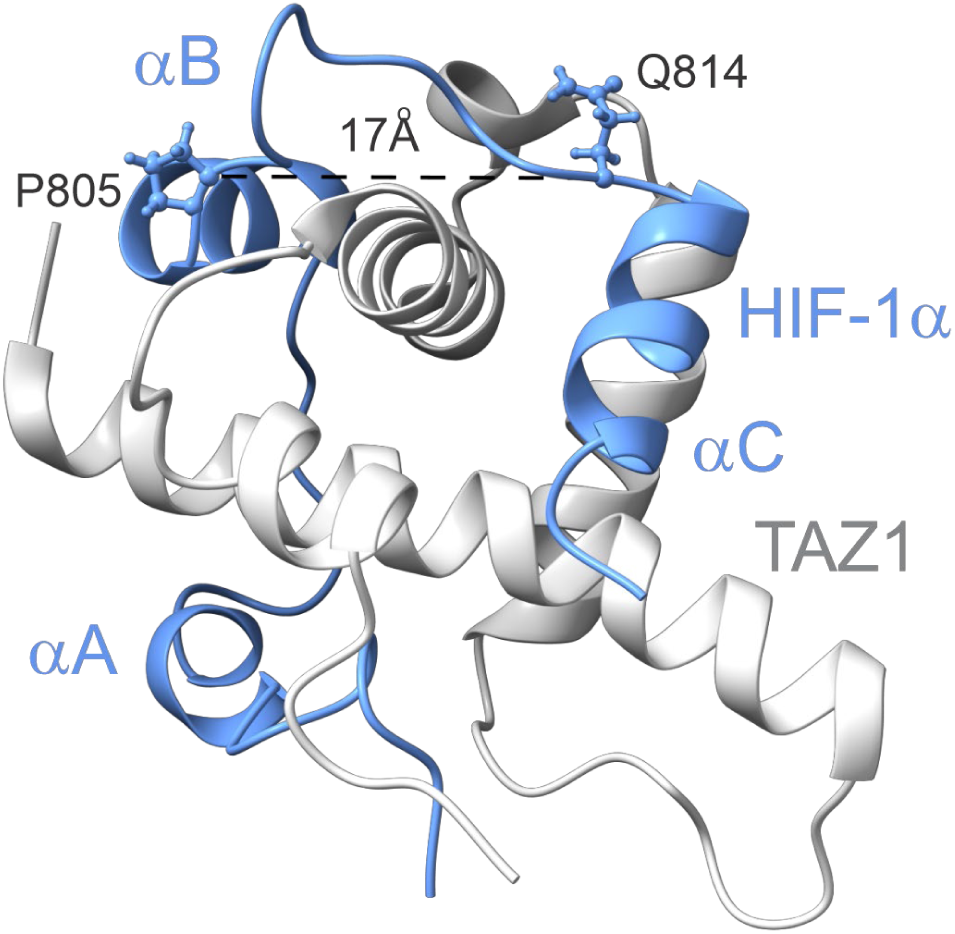
Solution structure of TAZ1:HIF-1α complex (PDB ID: 1L8C). The distance between the backbone Cα residues of P805 and Q814, at either end of the αB - αC linker, is indicated as a dashed black line.

The CITED2 activation domain displaces HIF-1α from TAZ1 by forming a transient ternary complex via its N-terminal region, which forms a helix (CITED2 αA) in the bound state.^*10*^ In addition to αA, CITED2 contains an LPEL motif and a hydrophobic cluster (ϕC) that function synergistically with the αA region to displace HIF-1α from TAZ1 and inhibit rebinding.^*16*^ Our previous work suggests that two molecular mechanisms contribute to the unidirectionality of HIF-1α displacement. Structural studies of TAZ1 bound to a CITED2-HIF-1α fusion peptide, a model of the ternary complex, revealed negative allostery between the binding sites for the CITED2 αA helix and HIF-1α helix αC, so that when CITED2 is bound, binding of the HIF-1α αC region is inhibited.^*17*^ In addition, we suggested that the presence of a highly flexible linker between the αB and αC binding motifs of HIF-1α gives rise to weak thermodynamic coupling between them, limiting binding cooperativity and strongly impairing the ability of HIF-1α to rebind to the CITED2:TAZ1 complex.^*10,15*^

To obtain detailed insights into the role of the αB - αC linker in modulating the TAZ1 binding affinity of the HIF-1α activation domain and its ability to compete with TAZ1-bound CITED2, constructs with varying linker length and sequence were generated by mutagenesis. TAZ1 binding by the mutants was monitored by native gel assays, NMR, and fluorescence anisotropy measurements. We hypothesized that reduction in linker length would result in stronger thermodynamic coupling between the HIF-1α interaction motifs, higher binding cooperativity, and an increased ability to compete with CITED2, whereas increased linker length would weaken coupling and reduce the cooperativity of the multivalent interaction. We find that both the length and composition of the linker are important. While Gly-Ser insertions decreased TAZ1 binding affinity to an extent proportional to the number of residues added, truncated and mutated linker sequences enhanced or inhibited binding in a context-dependent manner. HIF-1α linker variants that enhance TAZ1 binding affinity are more effective than wild type HIF-1α in displacing CITED2 from its TAZ1 complex. Our studies show that the length and composition of the αB - αC linker are critical for the fine-tuned allosteric regulation of the HIF-1α/CITED2 hypoxic switch and highlight that disordered linkers can function as more than simple entropic spacers between binding motifs.

## Materials and Methods

### Protein expression and purification

CBP TAZ1 (residues 340-439) was expressed in *E. coli* BL21 (DE3) (DNAY) and purified as previously described.^*10,18*^ HIF-1α CTAD (776-826) was expressed and purified as previously described.^*19*^ Uniform ^15^N labeling of HIF-1α constructs and TAZ1 for NMR experiments was performed by expression in minimal media containing ^15^N ammonium chloride (0.5 g l^−1^) and ^15^N ammonium sulfate (0.5 g l^−1^) as sole nitrogen sources. Uniform ^15^N, ^13^C labeling was achieved by adding ^13^C glucose as the sole carbon source in the minimal medium. Purified samples were exchanged into NMR buffer (20 mM Tris (pH 6.8), 50 mM NaCl, 2 mM DTT, and 5% D_2_O).

For native gel assays, HIF-1α (residues 776-826 (CTAD))and CITED2 (residues 216-269 (TAD)) activation domain constructs were expressed with N-terminal His_6_ and GB1 fusion tags and a thrombin cleavage site using a co-expression system, as previously described.^*19*^ Activation domain mutants were generated by PCR using established methods.^*20*^ Small scale 100 ml LB cultures were lysed by sonication in 5 ml of freshly-prepared 50 mM Tris pH 8, 1 M NaCl, 8 M urea, 5 mM DTT, and each lysis supernatant was batch-purified on 500μl cOmplete His-tag purification resin (Sigma-Aldrich) equilibrated with lysis buffer. After loading, the resin was washed with 20 CV lysis buffer, then 20 CV 20 mM Tris pH 8, 1 M NaCl, 5 mM imidazole. Bound protein was then eluted with 4 ml 20 mM Tris pH 8, 200 mM NaCl, 200 mM imidazole. Eluted His_6_GB1-tagged activation domains were exchanged into 20 mM Tris pH 8, 100 mM NaCl, 2 mM DTT using NAP25 desalting columns and further purified on 1ml Capto Q Impres pre-packed columns (Cytiva) using a gradient from 20 mM Tris pH 8 to 20 mM Tris pH 8, 250 mM NaCl. Purified proteins were concentrated to ∼200 μM, exchanged into assay buffer (20 mM Tris pH 8, 100 mM NaCl, 1 mM DTT) by desalting column and stored at −80C. Protein purity and identity were verified by SDS PAGE, reversed phase HPLC and MALDI mass spectrometry. Protein concentrations were determined by absorbance at 280 nm using calculated extinction coefficients.

### Native agarose gel competition assays

Protein stocks were diluted to 100 μM in assay buffer (20 mM Tris pH 8, 100 mM NaCl, 0.5 mM DTT) for native gels. Samples consisted of isolated proteins or mixtures of TAZ1 with equimolar activation domains at final 20 μM concentration of each protein, plus 2 μl of 5X sample loading buffer (100 mM Tris pH 8, 50% glycerol, 10 mM DTT), diluted with assay buffer to a final 10 μl volume. Samples were incubated at room temperature for 20 minutes to allow equilibration prior to loading on a 4 mm-thick 0.8% agarose gel poured in gel running buffer (25 mM Tris base, 192 mM glycine, final pH ∼8.3 not adjusted). The gels were run at a constant 50V for 1.5 hours, stained with Coomassie Brilliant Blue and protein bands were quantified with Image Lab software (Bio Rad). Samples were run at least in triplicate and on multiple gels; band intensity values were averaged.

### Fluorescence anisotropy

Fluorescence anisotropy measurements were carried out on an ISS-PC1 photon-counting steady-state fluorimeter at 20 °C in buffer containing 20 mM Tris (pH 6.8), 50 mM NaCl, 2 mM DTT. For competitive binding assays, 10 nM C800V HIF-1α labeled at the remaining C780 with AlexaFluor 594 was initially bound to 125 nM TAZ1. Unlabeled variants of HIF-1α were titrated into the preformed complex and data were recorded over a range of concentration of the competing HIF-1α variants. Data were fitted in GraphPad software using previously described binding equations.^*21,22*^ For CITED2 competition assays, a preformed complex of Alexa-594 labeled CITED2 (10 nM) and TAZ1 (125 nM) was titrated with WT HIF-1α or the ΔSR HIF-1α variant. Anisotropy measurements at high salt concentrations were made in 20 mM Tris (pH 6.8), 250 mM NaCl buffer with 2 mM DTT. For competition with R810A, an N-terminally fused GB1 R810A HIF-1α was used for the competition assays. For all anisotropy measurements, Alexa-labeled binary complexes were equilibrated for 15 minutes prior to the addition of competing ligand and data were collected after 30 minutes.

### NMR spectroscopy

NMR samples were prepared in buffer containing 20 mM Tris (pH 6.8), 50 mM NaCl, 2 mM DTT, and 5% D_2_O. Spectra were recorded at 25 °C on Bruker 700MHz and 800MHz spectrometers equipped with cryoprobes. NMR data were processed using NMRPipe ^*23*^ and analyzed using NMRFAM-SPARKY.^*24*^ For NMR samples of binary protein complexes, the unlabeled component was present in a 1.5-fold molar excess. Backbone resonance assignments for ^15^N-labeled ΔSR and ΔGS HIF-1α CTAD constructs bound to unlabeled TAZ1 were transferred from the HSQC spectrum of the WT complex. Standard triple resonance experiments were performed to resolve ambiguities and confirm cross peaks assignments for uniformly ^13^C, ^15^N-labeled (GS)_2_ HIF-1α CTAD bound to TAZ1. For HIF-1α/CITED2 competition experiments, the fraction of TAZ1 bound to each of the ligands was determined from cross peak volumes in the ^1^H-^15^N HSQC spectra of ^15^N-labeled TAZ1 in the HIF-1α and CITED2 complexes. Experimental errors were estimated from the Gaussian fit error of the peak volume obtained from SPARKY. Results are reported only for unambiguously assigned resonances that do not overlap in the ^1^H-^15^N HSQC spectrum and that have sufficient signal to noise (S/N>10) such that peak volumes can be reliably quantified.

### Amino acid sequence alignments

Over 170 sequences annotated as HIFα (comprising HIF-1α sequences from vertebrate species with both HIF-1α and HIF-2α proteins, and HIFα sequences from invertebrate species) from represented eukaryotic species were extracted from NCBI (https://blast.ncbi.nlm.nih.gov/Blast.cgi) and from published work.^*25–28*^ HIFα protein sequences, as well as the CITED2 and CBP protein sequences from the corresponding organisms, were aligned using Clustal Omega (https://www.ebi.ac.uk/jdispatcher/msa/clustalo),^*29*^ Guidance2 (https://taux.evolseq.net/guidance/), and MUSCLE (https://www.ebi.ac.uk/jdispatcher/msa/muscle) and analyzed for conservation of structural features and inter-motif linker length and composition. Alignments of representative sequences from each group were displayed with Jalview (Jalview.org); species names are abbreviated using the format *Homo sapiens* to HomSa. CBP alignments also include sequences from species lacking HIFα. A list of all sequence accession numbers used in this work is provided in supplementary Table S1.

### Modeling of CTAD/TAZ1 structures

Models of CTAD from various species bound to human TAZ1 were created with Alphafold3.^*30*^ Since the TAZ1 sequence is highly conserved, the human sequence was used for building the models for better consistency, but models built with homologous HIF-1α and TAZ1 species gave similar results.

## Results

### Sequence conservation

Since the HIF-1α/CITED2 hypoxic switch is so highly conserved in vertebrates, we reasoned that the characteristic structural features of the protein domains orchestrating this switch likely co-evolved in HIF-1α, CITED2, and CBP/P300. An extensive alignment of HIF-1α CTAD amino acid sequences is shown in Figure 2. The sequences in the upper group are from vertebrates and the cephalochordate lancelet, all of which contain both HIFα and CITED2. The invertebrates in the middle group contain HIFα and Factor Inhibiting HIF (FIH) but have no CITED2. Finally, the invertebrates in the bottom group retain HIFα but have neither FIH nor CITED2.

**Figure 2.**
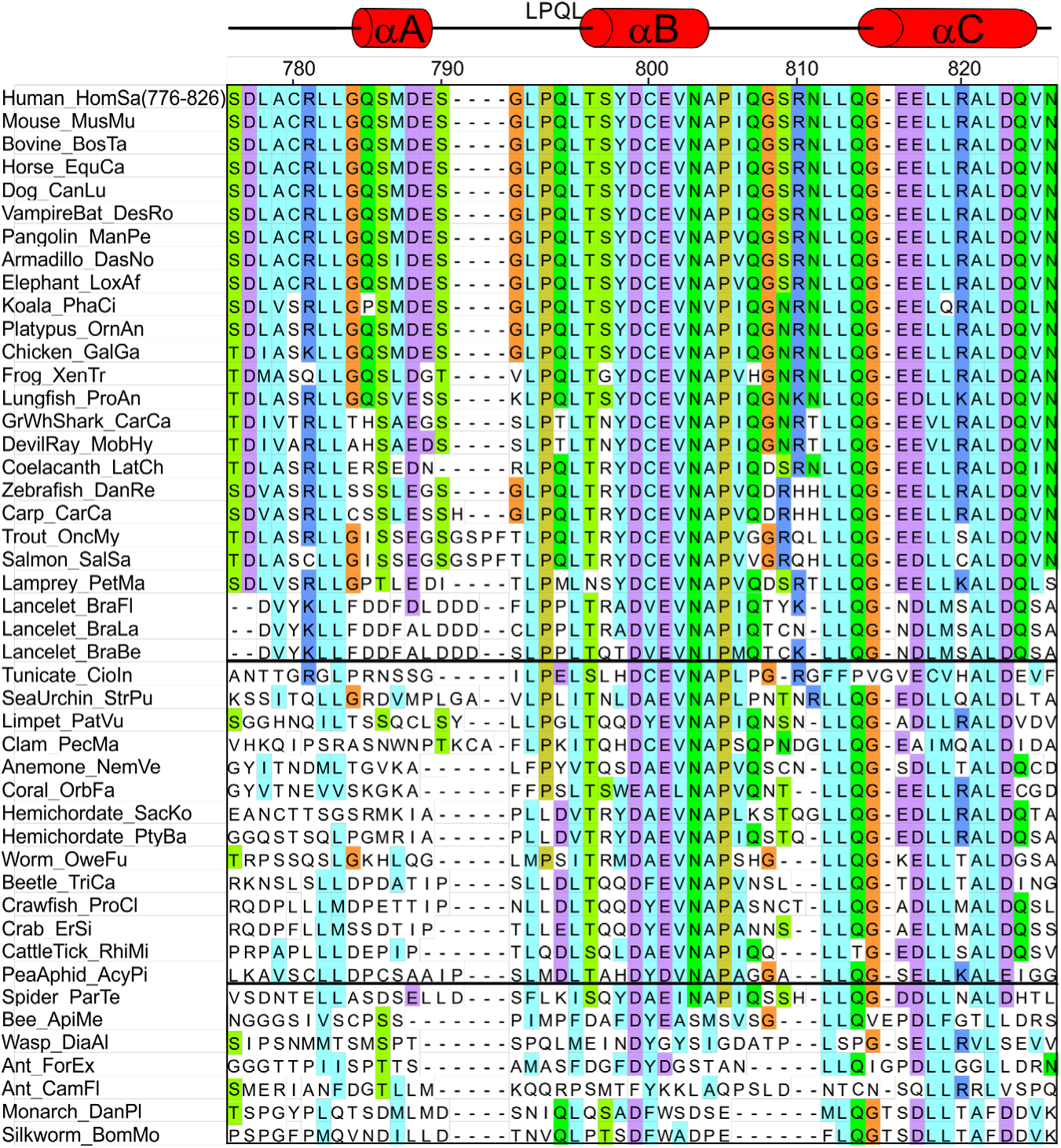
Alignment of CTAD sequences from representative vertebrate HIF-1α (upper section) and invertebrate HIFα (lower two sections) species. Secondary structure elements as defined previously^*16*^ are shown at the top of the figure. Numbers at the top of the figure refer to the human sequence.

The LPQL, αB and αC helical regions are highly conserved in vertebrates and the cephalochordate lancelet. Sequence conservation in the αA region and the αB - αC linker (residues 805-811) is confined to the higher vertebrates and both regions vary in composition and length in fish and lancelet. While the sequences of the αB and αC regions are largely conserved in invertebrates that contain FIH, the sequence of the αB - αC linker is highly variable, likely reflecting the absence of CITED2 and, as a consequence, of the HIF/CITED regulatory switch. The amino acid sequences of CITED2 are all highly conserved (Figure S1A), arguing for a common molecular mechanism for the unidirectional switch by which CITED2 regulates HIF-1α.^*10*^ The CBP TAZ1 domain interacts with many transcription factors and its sequence is highly conserved even in species lacking HIF-1α and CITED2 (Figure S1B).

The conservation of HIF-1α and HIFα CTAD sequences is an indication that interaction with TAZ1 is likely to be structurally similar, especially in the αB and αC regions. Alphafold3 models of bound HIF-1α CTADs from orthologs show the highly conserved LPQL-αB - αC cassette docked at the same TAZ1 interface as the human HIF-1α CTAD (Figure S2), even for invertebrate species with short αB - αC linkers and altered LPQL motifs like coral (FPSL) and beetle (LLDL). For many orthologs, helical structure in the αA region is predicted with low confidence and this region is often disordered or not docked onto TAZ1 in AlphaFold 3 models.

Interestingly, the linker between the C-terminus of αB and the N-terminus of αC, which is disordered through much of its length in the TAZ1-bound state,^*11,12,15*^ is highly conserved in length and composition in vertebrates. In mammals and reptiles, the sequence in the central region of the linker, QG(S/N)RN, is highly conserved but it begins to diverge in fish, coelacanth and lamprey. In lancelet and in invertebrate species lacking CITED2, the αB - αC linker is highly divergent in composition and is mostly truncated by deletion of 1 to 3 residues (Figure 2). The contrast between the conservation of the linker between species that do or do not contain CITED2, together with the observation that thermodynamic linkage between αB and αC of HIF-1α is weak,^*10*^ led us to hypothesize that the length and composition of the linker may be a decisive factor in regulating the HIF-1α-TAZ1-CITED2 unidirectional switch.

### Screening of variants by native gel electrophoresis

Mutants of the HIF-1α CTAD were designed to vary the length, flexibility, and amino acid composition of the linker. To facilitate mutagenic screens, we developed a native gel competition assay to measure relative affinities of bacterially-expressed activator and coactivator domain constructs. Activation domain constructs purified at small scale as His_6_GB1 fusions were used directly in the assay without cleaving the fusion tag, which allows for accurate concentration determination by UV absorbance and better detection with Coomassie stain. Relative affinities were measured by competition with WT CTAD without the fusion tag. TAZ1 complexes with HIF-1α CTAD with and without the His_6_GB1 fusion tag have different migration patterns on native gels, so mixtures of TAZ1 complexes of mutant HIF-1α CTAD (with fusion tag) and WT HIF-1α CTAD (without fusion tag) can be resolved on the gel, and the intensities of bands representing the HIF-1α CTAD fusion/TAZ1 complex can be measured after Coomassie staining. This technique gives a quick, semi-quantitative measure of relative affinity changes and reproduces trends observed in solution NMR and fluorescence anisotropy assays without having to label the proteins with isotopes or fluorescent dyes. An overview of the method and a representative gel are shown in Figure 3 and results of competition screening experiments between His_6_GB1-HIF-1α CTAD mutants (ligand 1 or L1) and WT HIF-1α CTAD (ligand 2 or L2) are summarized in Table 1.

**Figure 3.**
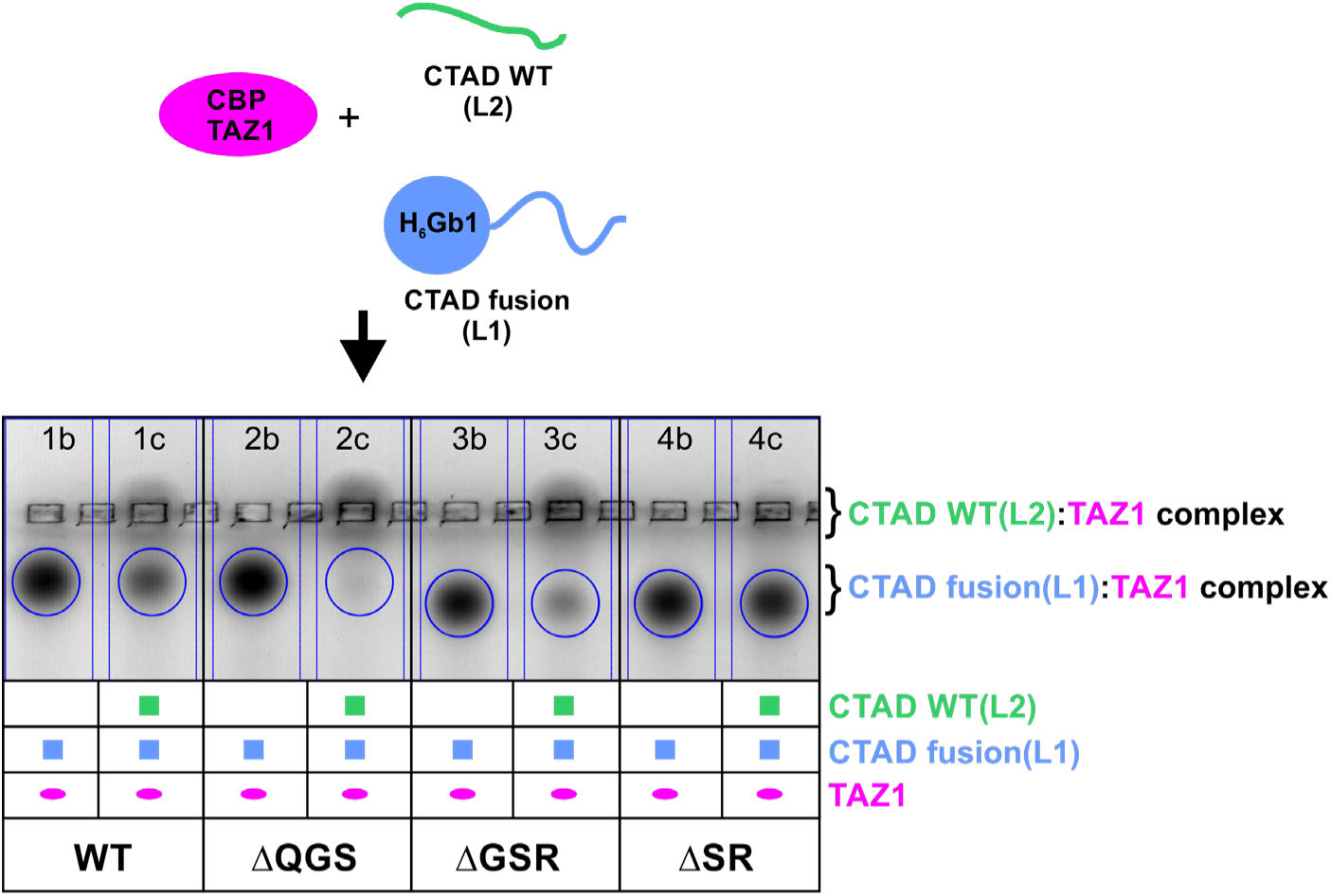
Schematic diagram of native gel competition assay and representative gel showing competition of His_6_GB1-HIF-1α CTAD fusion (Ligand 1 or L1) with tag-free WT HIF-1α CTAD (Ligand2 or L2) for binding to TAZ1. Each pair of lanes contains the His_6_GB1-HIF-1α CTAD (L1) constructs as annotated in the last row below the gel. Binary (b) lanes contain TAZ1 and L1 at 1:1 molar ratio. Competition (c) lanes contain TAZ1, L1, and L2 at 1:1:1. Labels at the bottom of the figure show which L1 His_6_GB1 fusion (WT HIF-1α CTAD or mutants identified in Table 1) is in competition with tag-free WT HIF-1α CTAD. Note that complexes of tag-free CTAD with TAZ1 stay at the position of the injection well. Circles around the His_6_GB1-HIF-1α CTAD (L1):TAZ1 complex band denote regions used for intensity determination.

**Table 1.**
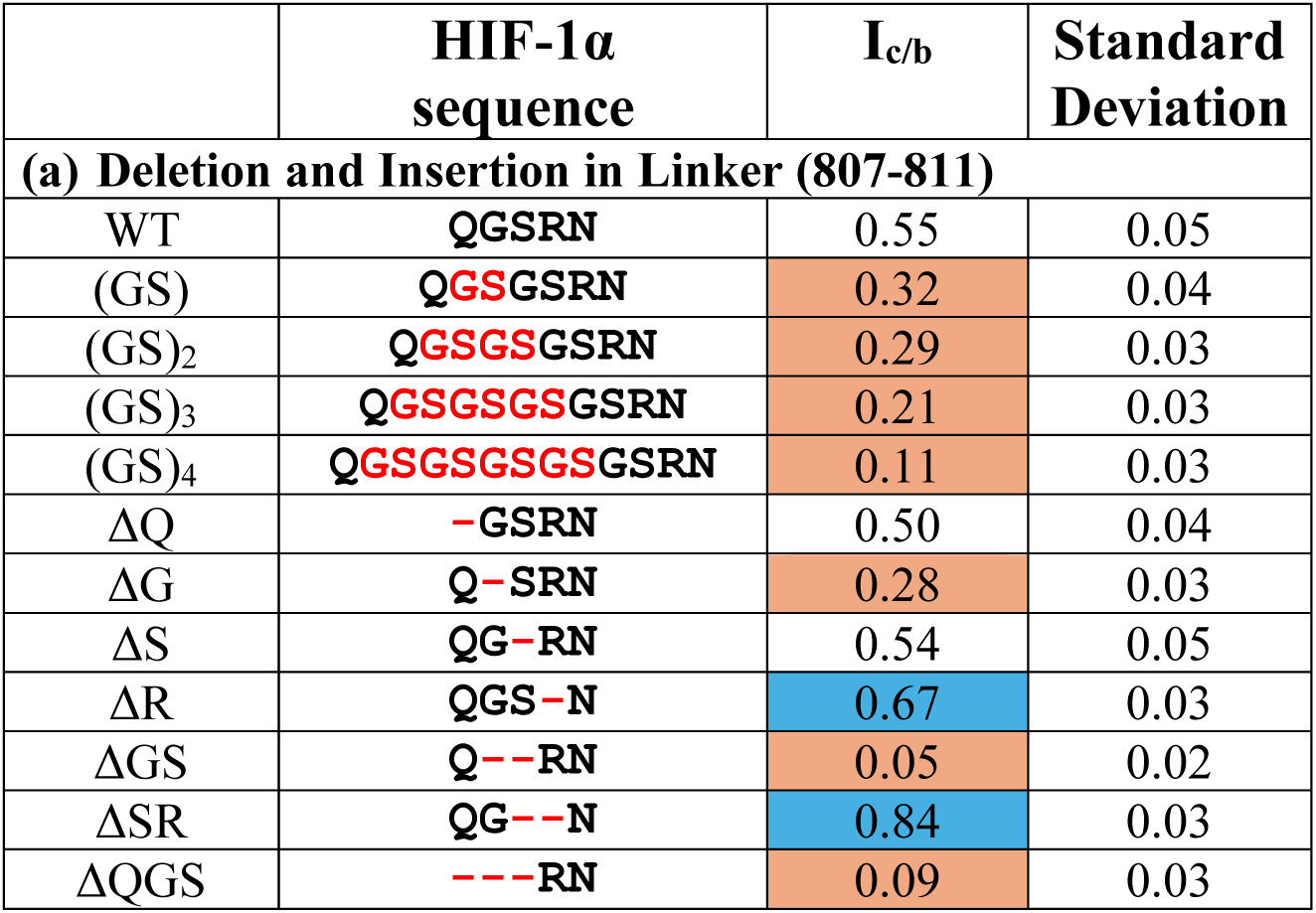
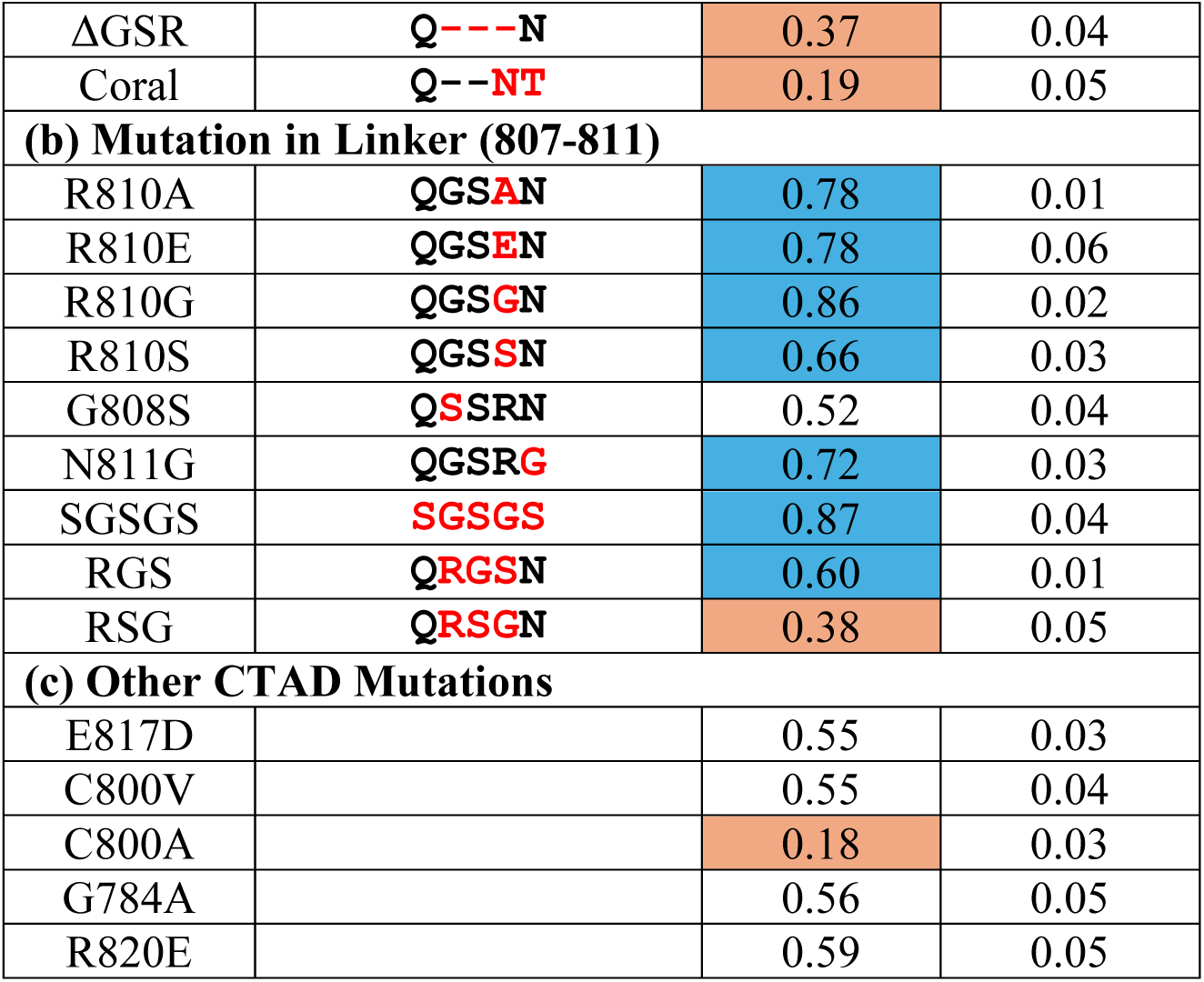
Relative TAZ1 affinities of HIF-1α CTAD mutants derived from native gel competition assay screens. The relative affinity of mutant HIF-1α CTAD relative to WT for TAZ1 is described by the I_c/b_ ratio, which represents relative populations of the TAZ1 complexes in native gel competition assays as shown in Figure 3. The HIF-1α CTAD constructs are grouped by mutation type: (a) linker insertions and deletions, (b) linker mutations, and (c) mutations outside the αB - αC linker. Mutations that increase the affinity (I_c/b_ > 0.6) are highlighted in cyan and those that decrease the affinity (I_c/b_ < 0.4) in orange.

The gel in Figure 3 shows pairs of assays, with (c = competition) and without (b = binary) competition by tag-free WT HIF-1α CTAD. The ratio of the intensities (I_c/b_) of the TAZ1/L1 band (circled) in the competition lane relative to the binary lane is proportional to the relative affinities of mutant HIF-1α (L1) to WT HIF-1α (L2) CTAD. If the affinities of L1 and L2 are equal, the intensity in the c lane (TAZ1:L1:L2 = 1:1:1) should be 0.5 times that in the b lane (TAZ1:L1 = 1:1). A HIF-1α CTAD mutant that binds more tightly to TAZ1 than WT CTAD will have I_c/b_ greater than 0.5 while, conversely, a mutant that binds TAZ1 more weakly will have I_c/b_ less than 0.5. Lanes 1b and 1c show the control experiment for competition between WT HIF-1α CTAD with and without the fusion tag (L1 and L2, respectively), where TAZ1 is fully bound to L1 in the binary lane but only ∼50% bound to L1 in the competition lane and I_c/b_ = 0.55 ± 0.05 (Table 1), demonstrating that the His_6_GB1 tag does not interfere with TAZ1 binding or competition between the two HIF-1α CTAD constructs. Control experiments were also done with the HIF-1α CTAD C800V mutant used for fluorescent dye labeling and anisotropy assays, which has similar affinity to the wild type, and with the C800A mutant, which has decreased affinity.^*16,31*^ In the native gel competition assay, C800V has I_c/b_ = 0.55 ± 0.04, while C800A has I_c/b_ = 0.18 ± 0.03, which is consistent with prior measurements. Additionally, a native gel assay of competition for TAZ1 binding between HIF-1α and CITED2 CTADs (Figure S3) shows that in a 1:1:1 mixture of TAZ1 with HIF-1α and CITED2, TAZ1 is fully bound to CITED2 CTAD, as previously observed in NMR studies.^*10*^ Thus, even though this is a non-equilibrium binding assay, the native gel method reproduces solution binding behavior for the high affinity (K_d_ = 10 nM) TAZ1/HIF-1α system.

If the hypothesis that the disordered linker between αB and αC of HIF-1α is important for the operation of the unidirectional switch is correct, we expect that increasing the length of the linker by insertion of Gly-Ser dipeptide repeats should further decouple the αB and αC motifs and decrease the affinity for TAZ1. This is what we observe (Table 1). Decreasing the length of the linker, which conversely should enhance the affinity by reducing conformational freedom, proved to be informative since these mutations revealed residue-specific effects. Structural models generated using AlphaFold 3^*30*^ confirm that as many as three residues can be deleted from the central region of the linker without affecting interactions of the αB and αC helices with TAZ1. Mutants with single amino acid deletions of linker residues Q807 and S809 have similar affinity to the wild type CTAD, but deletion of G808 or R810 has a significant and opposite effect on the affinity, with ΔG808 binding TAZ1 more weakly (I_c/b_ = 0.28) than WT and ΔR810 binding more tightly (I_c/b_ = 0.67). Deletion of two linker residues in the ΔSR(809-810) mutant further enhances the affinity (I_c/b_ = 0.84). Substitution of R810 with Ala, Gly, Ser or Glu also enhances the binding affinity, underscoring the importance of electrostatic effects in interactions between the HIF-1α CTAD and the electropositive TAZ1 domain.^*32*^ Mutation of R820, on the αC helix, to Glu only moderately increases the affinity (I_c/b_ =0.59), indicating that tighter binding of R810 mutants is not solely a result of decreasing the overall HIF-1α CTAD positive charge. Interestingly, R810 is poorly conserved in species lacking CITED2 (Figure 2).

In contrast, deletion of two linker residues in the ΔGS(808-809) mutant results in even weaker binding to TAZ1 (I_c/b_ = 0.05) than the single deletion mutant ΔG808 (I_c/b_ = 0.28) and points to a requirement for a flexible hinge in the linker for optimal binding. Deletion of three linker residues in the ΔQGS(807-809) or ΔGSR(808-810) mutants results in significant loss of TAZ1 binding affinity (I_c/b_ = 0.09 and 0.37, respectively). The effects of the deletion mutations seem to be additive, with deletion of R810 in ΔGSR partially compensating for affinity loss from deleting three linker residues. Mutant Coral, in which linker residues GSRN(808-811) are replaced with the sequence from *Orbicella faveolata* (NT), resulting in a two-residue deletion, has reduced binding in native gel competition assays (I_c/b_ = 0.19), consistent with the combined effects of mutating R810 and deleting GS(808-809).

The requirement for flexibility in the αB - αC linker was further studied with mutants G808S, N811G, and SGSGS, and with the scrambled mutants RGS and RSG. Unlike the G808 deletion mutant, the G808S mutant retains wild type affinity, consistent with a general requirement for flexibility in this region when the linker length is unchanged. Mutants RGS, RSG and N811G have linkers with the same overall charge and length as the wild type CTAD, but RSG has decreased affinity (I_c/b_ = 0.38) and RGS has a small enhancement in affinity (I_c/b_ = 0.60). Mutant N811G, which incorporates a second Gly residue in the linker, has further enhanced affinity (I_c/b_ = 0.72). Mutant SGSGS has the highest affinity of all CTAD variants tested (I_c/b_ = 0.87), consistent with the combined effects of decreasing positive charge at R810 and enhancing linker flexibility. Overall, these results demonstrate stringent requirements on linker length, charge and relative placement of segments with enhanced flexibility.

### Effect of enhancing HIF-1α CTAD affinity for TAZ1 on competition with CITED2

Competition assays monitored by native gel electrophoresis also show that enhanced-affinity HIF-1α CTAD mutants are more effective than wild type HIF-1α at competing with CITED2 TAD for binding to TAZ1 (Figure S3). Equimolar mixtures of TAZ1 with CITED2 and the higher affinity HIF-1α CTAD mutants contain predominantly TAZ1:CITED2 complex, but when high affinity HIF-1α CTAD mutants are added at up to four-fold excess, a band corresponding to the TAZ1:HIF-1α complex is observed for the mutants but not for wild type HIF-1α CTAD at four-fold excess. The deletion mutant ΔSR, the variant in which R810 is mutated to G (keeping the linker length unchanged), and the variant in which the native QGSRN linker is replaced by SGSGS, all show increased affinity for TAZ1 and enhanced ability to compete with CITED2 in the native-gel assays.

### Measurement of K_d_ for HIF-1α variants using fluorescence anisotropy

To gain a more precise understanding of the effect of changes in the αB - αC linker, we measured the affinity with which several of the HIF-1α variants bind TAZ1 using fluorescence anisotropy competition experiments.^*21,22*^ We selected HIF-1α variants with reduced linker lengths that exhibited the largest difference from WT in native-gel binding assays, as well as the series of GS repeat insertions to probe the effect of gradual lengthening of the intervening linker (Table 1). The variant HIF-1α constructs were titrated into a preformed complex of TAZ1 with C800V HIF-1α CTAD labeled at C780 with AlexaFluor 594. The apparent K_d_ determined for WT HIF-1α CTAD (K_d_ =14 ± 2 nM) is close to the K_d_ of the binary HIF-1α:TAZ1 complex reported previously (K_d_ = 10 ± 1 nM).^*10*^ Consistent with the gel binding assays, the gradual increase of the linker length by insertion of multiple Gly-Ser repeats (Q(GS)_n_GSRN) reduces the binding affinity from 5 to 12-fold for n = 1 to 4 (Figure 4, Table 2). These data highlight the sensitivity of the gel assays; a modest 5-fold increase in K_d_ for a single GS insert is reflected in a change in I_c/b_ from 0.55 to 0.32 (Figure 4, Table 1).

**Figure 4.**
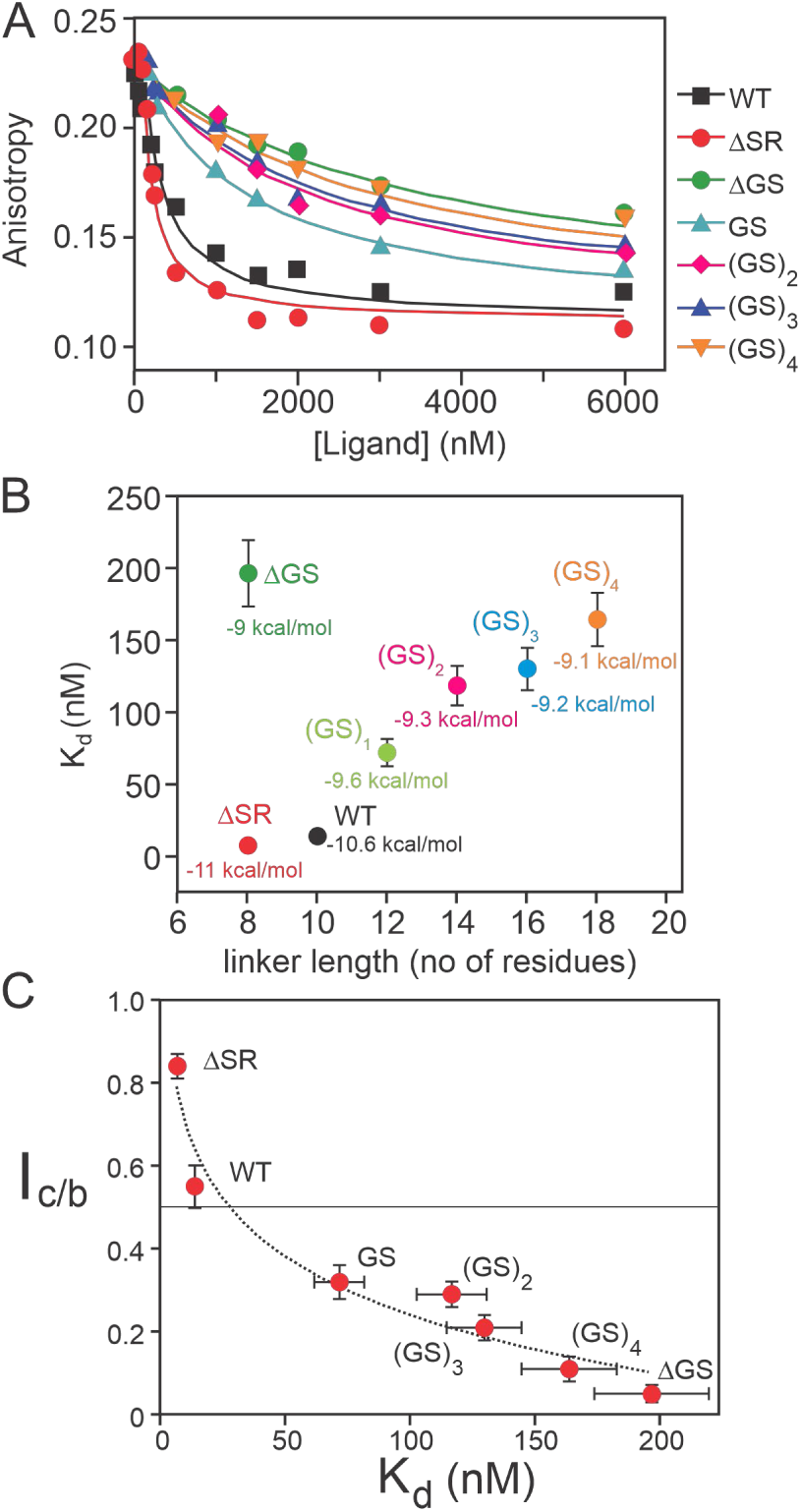
Fluorescence anisotropy competition assay. Unlabeled HIF variants were titrated into a preformed complex of AlexaFluor-594 labeled WT HIF-1α peptide and unlabeled TAZ1. (a) Fits of the anisotropy competition assay for respective HIF variants. (b) TAZ1 binding affinity of HIF-1α variants plotted as a function of intervening linker length. Corresponding binding free energy (at 293K) is labeled in the plot. (c) Plot of I_c/b_ vs K_d_ from fluorescence anisotropy with logarithmic trend line.

**Table 2.** Comparison of K_d_ values determined by fluorescence anisotropy competition experiments and I_c/b_ values from native agarose gel competition assays. Uncertainties in K_d_ values are fitting errors. Reported I_c/b_ values are the average and standard deviation from independent gel competition assays (n ≥ 3).

| Ligand | $K_d$ (nM) | $I_{c/b}$ |
| --- | --- | --- |
| WT | $14 \pm 2$ | $0.55 \pm 0.05$ |
| ΔSR | $7 \pm 1$ | $0.84 \pm 0.03$ |
| ΔGS | $200 \pm 20$ | $0.05 \pm 0.02$ |
| (GS) | $70 \pm 10$ | $0.32 \pm 0.04$ |
| (GS)<sub>2</sub> | $120 \pm 10$ | $0.29 \pm 0.03$ |
| (GS)<sub>3</sub> | $130 \pm 15$ | $0.21 \pm 0.03$ |
| (GS)<sub>4</sub> | $160 \pm 20$ | $0.11 \pm 0.03$ |

Consistent with the results from the agarose gel competition experiments, we found that shortening of the native linker has very different effects on binding depending on the nature of the deletion. The ΔSR-HIF-1α construct forms the highest affinity (K_d_ = 7 ± 1 nM) binary TAZ1 complex while deletion of the GS(808-809) dipeptide resulted in the weakest affinity (K_d_ = 200 ± 20 nM, I_c/b_ = 0.05) among the HIF-1α variants tested.

### Effect of linker mutations on interactions of the individual binding motifs with TAZ1

Structural changes in the binary complexes of TAZ1 with HIF-1α CTAD constructs with variant linkers could potentially be responsible for the observed changes in binding affinity. To explore this possibility, we characterized the binary HIF-1α/TAZ1 complexes using solution NMR spectroscopy. Three variants were selected for NMR analysis, ΔSR HIF-1α, (GS)_2_ HIF-1α, and ΔGS HIF-1α, which show a wide range of TAZ1 binding affinities. Comparison of ^1^H-^15^N heteronuclear single quantum coherence (HSQC) spectra of the unbound CTAD variants (Figure S4) shows no difference in the backbone amide chemical shifts except at the site of the mutated αB - αC linker. In particular, the complete overlap of the cross peaks associated with the αB, αC, and LPQL motifs shows that the changes in the linker sequences do not affect the intrinsic conformational propensities of the HIF-1α binding motifs and that the peptides remain disordered. The HIF-1α CTAD folds upon binding to TAZ1, resulting in large changes in the HIF-1α HSQC spectrum.^*11*^ Weighted average ^1^H, ^15^N chemical shift differences between the free and TAZ1-bound state are very similar for WT HIF-1α and the ΔSR, (GS)_2_, and ΔGS variants (Figure 5), showing that all undergo the same structural changes upon binding to TAZ1. These observations indicate that the changes that we observe in binding affinity arise from changes in the length and composition of the linker, rather than differences in the interaction of the individual LPQL, αB and αC binding motifs with TAZ1 or differences in the structures or conformational propensities of the unbound and TAZ1-bound HIF-1α CTAD.

**Figure 5.**
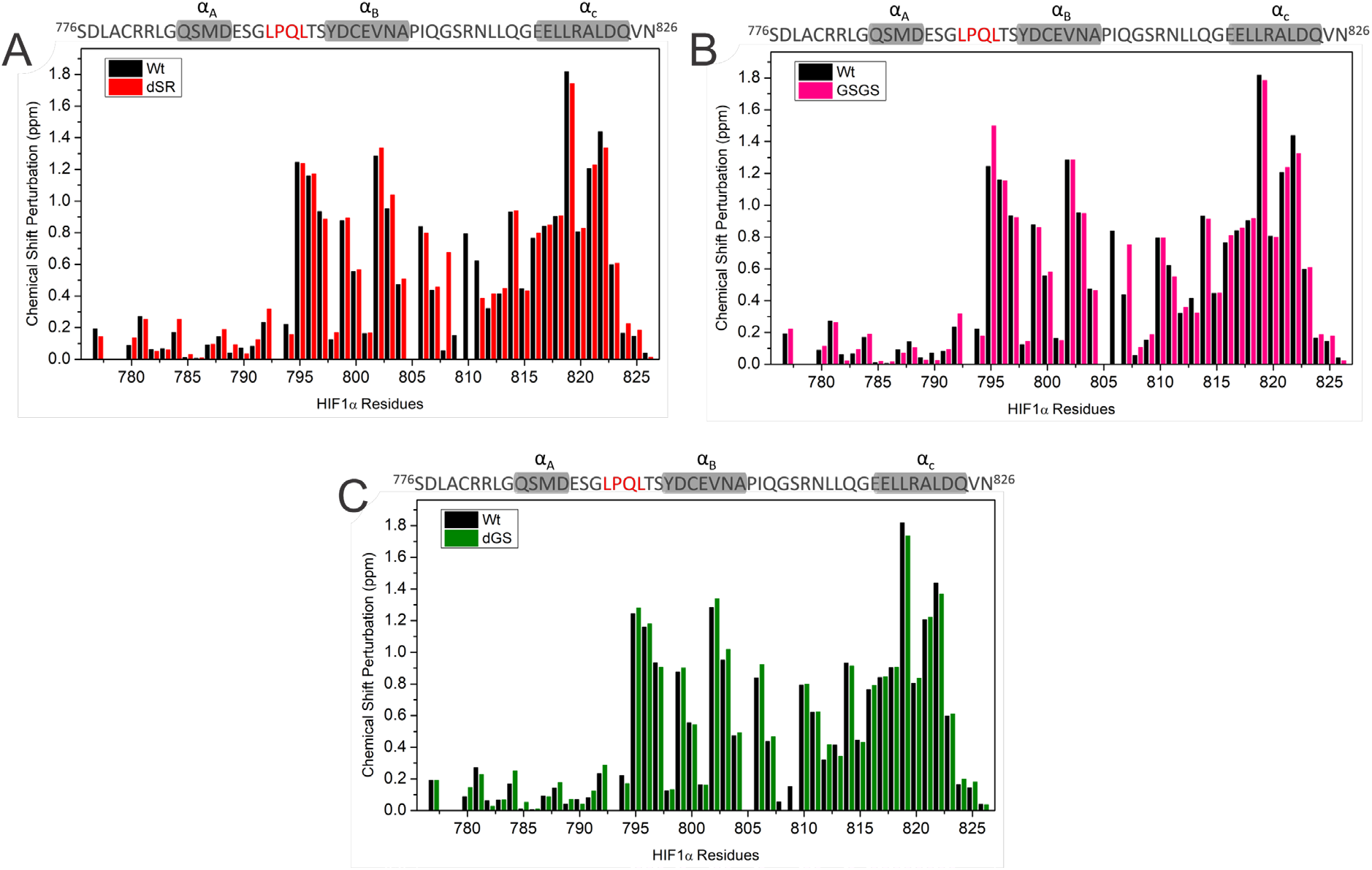
Comparison of weighted average ^1^H, ^15^N chemical shift differences (Δδ_ave_ = [(Δδ_HN_)^2^+(Δδ_N_/5)^2^]^1/2^) between free and TAZ1-bound state WT HIF-1α (black) and the variants (a) ΔSR (red), (b) (GS)_2_ (magenta), and (c) ΔGS (green).

### HIF-1α linker modifications perturb the unidirectional allosteric switch

The negative regulator CITED2 displaces the HIF-1α CTAD from TAZ1 employing a unidirectional switch mechanism that was characterized by fluorescence anisotropy and NMR.^*10*^ CITED2 displaces HIF-1α very efficiently (apparent K_d_ ∼ 0.2 nM), whereas HIF-1α is much less efficient at displacing TAZ1-bound CITED2 (apparent K_d_ ∼ 900 nM). As a follow-up to the native gel assays (Figure S3), we performed fluorescence anisotropy and NMR experiments to investigate the effect of αB - αC linker mutations that increase the TAZ1 binding affinity on the reversibility of the switch. Fluorescence anisotropy competition experiments in which WT HIF-1α CTAD or ΔSR HIF-1α CTAD were titrated into an Alexa594-CITED2/TAZ1 complex showed that the higher-affinity ΔSR HIF-1α variant is much more efficient in displacing bound CITED2 (apparent K_d_ = 85 ± 9 nM for ΔSR HIF-1α versus 1100 ± 200 nM for WT HIF-1α, Figure 6).

**Figure 6.**
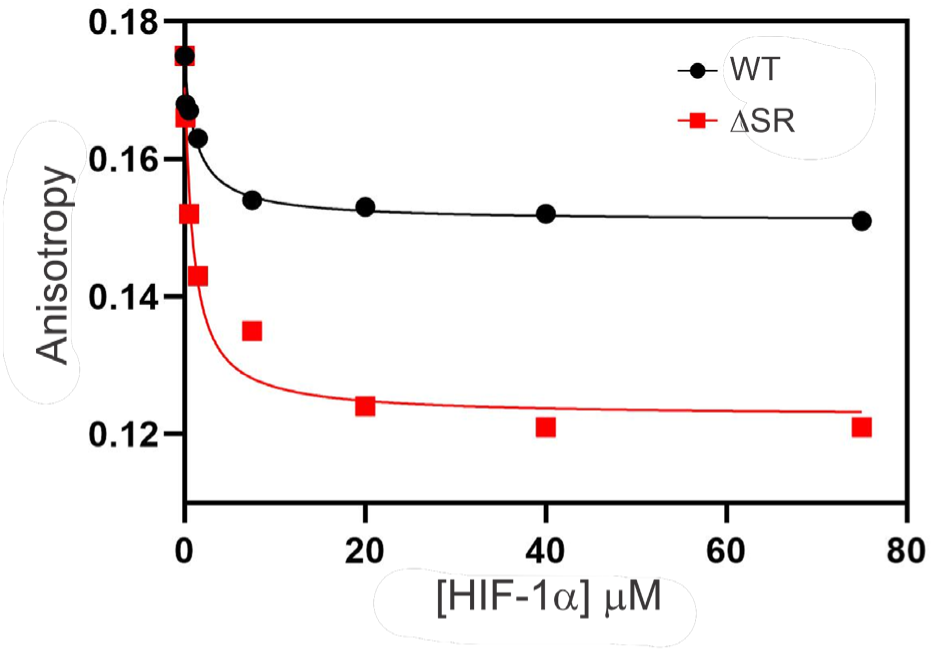
Fluorescence anisotropy competition assay. A pre-formed complex of unlabeled TAZ1 with CITED2 labeled with Alexa-Fluor 594 was mixed with increasing concentrations of WT (black) or ΔSR mutant (red) HIF-1α CTAD.

The ability of ΔSR HIF-1α to compete effectively with CITED2 was also examined by NMR. Many amide cross peaks in the ^1^H-^15^N HSQC spectra of ^15^N-labeled TAZ1 have distinct chemical shifts in the HIF-1α and CITED2 complexes HIF-1α (see Figure S5).^*10,16*^ Measurement of peak volumes or intensities of corresponding cross peaks from different complexes enables a quantitative measure of the relative population of TAZ1:HIF-1α and TAZ1:CITED2 complexes when both ligands are present. Two examples are shown in Figure 7. HSQC spectra of ^15^N-labeled TAZ1 mixed with WT HIF-1α and CITED2 CTAD peptides in a 1:1:1 molar ratio contain only cross peaks corresponding to the TAZ1:CITED2 complex.^*10*^ In contrast, the HSQC spectrum of an equimolar mixture of ^15^N-TAZ1, CITED2, and ΔSR HIF-1α shows a 16% population of ΔSR HIF-1α -bound peaks. Increasing the molar fraction of ΔSR HIF-1α to 2:1 and 4:1 relative to CITED2 further increases the fraction of TAZ1-bound ΔSR HIF-1α to ∼40% (Figure 7), compared to a population of ∼0.05% for the corresponding mixture containing WT HIF-1α. A superposition of the full ^15^N-TAZ1 ^1^H-^15^N HSQC spectra is shown in Figure S6.

**Figure 7.**
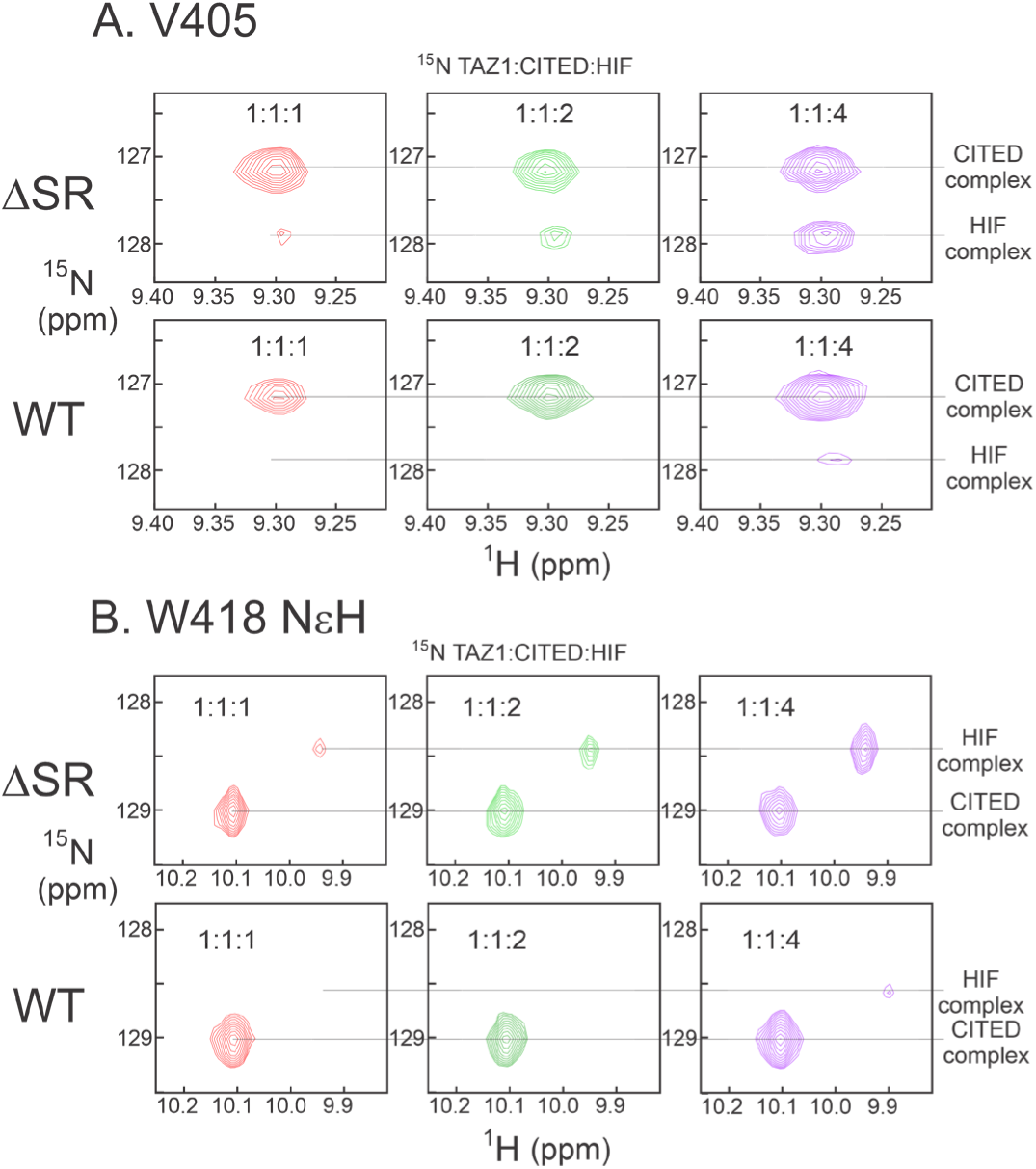
Regions of ^1^H-^15^N HSQC spectra of ^15^N-labeled TAZ1. A. Region showing the cross peak of V405 NH in complex with CITED2 and HIF-1α TAD ΔSR (top row) and WT (second row) in the ratios 1:1:1 (red), 1:1:2 (green) and 1:1:4 (purple). B. Region showing the cross peak of W418 NεH in complex with CITED2 and HIF-1α TAD ΔSR (third row) and WT (fourth row).

## Discussion

A robust response to low oxygen tension (hypoxia) is absolutely required for obligately aerobic organisms. Vertebrate animals have evolved an exquisitely tuned hypoxic response that employs the versatile capabilities of intrinsically disordered proteins to control and mediate the process. The hypoxia-inducible factor (HIF) contains both structured and disordered domains; the structured domains include per-arnt-sim (PAS) domains concerned with protein-protein interactions. The disordered domains include the ODD domain that mediates the proline hydroxylation that targets HIF for proteasomal degradation under normoxic conditions, through interaction with the von Hippel-Lindau factor.^*6*^ The disordered C-terminal activation domain contains a conserved asparagine N803 that is also hydroxylated, by an enzyme termed factor inhibiting HIF, or FIH.^*33*^ Under normoxic conditions, HIF is inactivated through these hydroxylation reactions. Under hypoxic conditions, HIF is not hydroxylated at proline or N803, is stabilized against degradation and activates transcription of hypoxia-responsive genes by interaction with the TAZ1 domain of the transcriptional coactivators CBP and p300.^*7*^ When hypoxic conditions are ameliorated, the hypoxic response must be rapidly and completely down-regulated. This is achieved through the action of another disordered protein, CITED2, one of the downstream genes transcribed in response to HIF which functions as a negative feedback regulator.^*9*^ The molecular mechanism of this response illustrates the exquisite control of these processes by intrinsically disordered protein domains.

### Sequence conservation of the hypoxia pathway

The TAZ1-HIF-1α-FIH-CITED2 system is highly conserved in vertebrates, and patterns of sequence conservation in the protein components suggest that they have co-evolved. The CITED2 TAD sequence is invariant in vertebrates (Figure S1A) and differences are only observed in cephalochordates (lancelet), which are at the dividing line between the organisms with and without the hypoxic switch. Placozoans, the earliest animals, have a HIFα gene but lack CTAD and FIH (as well as CITED2), consistent with the presence of CTAD in more complex organisms requiring more nuanced regulation of the hypoxic response.^*34*^ A correlation between the conservation of HIF N803 and the presence in an organism’s genome of the enzyme FIH, which mediates N803 hydroxylation, has been noted previously.^*34*^ It appears that sequence conservation in the αA region of HIF and in the linker between αB and αC correlates with the presence in the organism of CITED2, an indication that the unidirectional switch that mediates the shutoff of the hypoxic response is exclusive to higher-order organisms such as mammals and birds.

### Effect of linker length on the unidirectional switch

Upon binding to the TAZ1 domains of CBP and p300, HIF-1α folds to form helices αB and αC connected by a 10-residue linker (P805-Q814). The central region of the linker (Q808 – L812) is disordered in the NMR structures of the complex,^*11*^ (Figure 8) and ^15^N relaxation measurements show that the linker backbone is highly dynamic, with ^15^N order parameters (S^2^) < 0.5 for all residues except N811, indicating large amplitude backbone fluctuations.^*15*^ Consistent with this, residues QGSRNL have elevated backbone B-factors in HIF-CITED fusion proteins bound to the CBP (PDB ID 7LVS) and p300 (7QGS) TAZ1 domains.^*17,35*^ The ends of the linker are anchored by sidechain interactions of the hydrophobic residues I806 and L813, which pack against the TAZ1 surface in the NMR structures of the CBP TAZ1:HIF-1α complex.^*11*^ The γ_2_ methyl group of I806 is motionally restricted (S^2^ = 0.78), whereas the δ_2_ methyl group of L812, within the linker, has a below-average S^2^ (0.36).^*36*^

**Figure 8.**
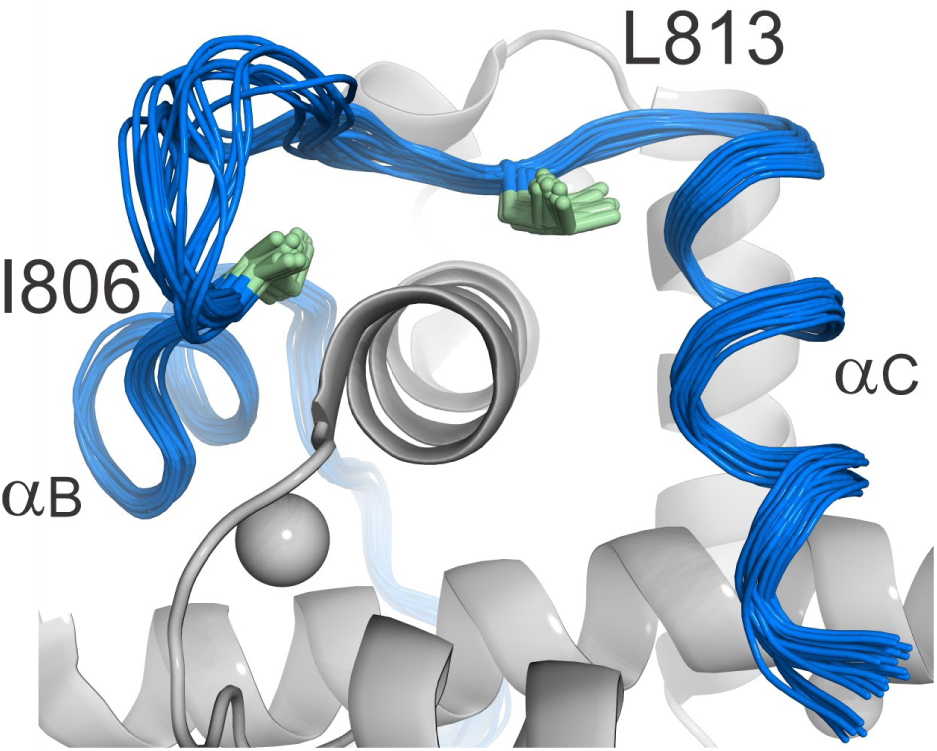
Region of the NMR structure family^*11*^ of the complex between the CTAD of HIF-1α (blue) and TAZ1 (gray), showing the disorder in the loop between αB and αC and the well-ordered side chains of I806 and L813 at either end of the disordered loop.

Increasing the length and flexibility of the αB - αC linker by insertion of a series of GS dipeptides causes a monotonic decrease in the affinity (avidity) of the multivalent HIF-1α CTAD for TAZ1 (Table 1, Figure 4). The decrease in avidity results from an increase in the entropic cost of binding as the length of the flexible linker is increased, decreasing the effective concentration and weakening the thermodynamic coupling between the αB and αC binding motifs.^*37,38*^ These results explain the invariant linker length in vertebrates; insertion of even a single GS dipeptide decreases the affinity by 5-fold, suggesting that any increase in linker length would likely impair the hypoxic response.

The results obtained with the linker deletions and the positional and scrambled mutants indicate that the composition of the linker is also highly important. R810 is associated with a consistent decrease in the affinity of the HIF-1α CTAD for TAZ1 and arginine is strictly conserved at this position in tetrapods. The linker sequence in fish is more variable but all contain an arginine residue near the middle of the linker. The NMR structures of the HIF-1α-TAZ1 complex^*11*^ and the X-ray crystal structure of the TAZ1:fusion protein complex^*17*^ show that R810 is close to an electropositive region of the TAZ1 surface and the positive end of the protein dipole (dipole moment 380 Debye, calculated from the X-ray structure using the Protein Dipole Moments server^*39*^). Single-site substitution of R810 by A, E or S to remove the arginine charge relieves unfavorable interactions with TAZ1 and enhances HIF binding affinity (I_c/b_ = 0.66 – 0.78 compared to 0.55 for WT). Substitution of R810 by G further increases affinity (I_c/b_ = 0.86), probably due to increased flexibility of the linker.

Scrambling of the central region of the linker sequence from GSR to RGS has only a small effect, increasing the affinity slightly (I_c/b_ increases from 0.55 to 0.60). In contrast, swapping G808 and R810 to change the central sequence to RSG impairs binding, with I_c/b_ decreasing to 0.38. This result is initially surprising, given the substantial increase in affinity of the single site mutation R810G. However, the R810G mutant contains two Gly residues in the linker (QGSGN), which will thus be inherently more flexible than the WT QGSRN linker. The stiffness of a disordered linker depends strongly on the presence of Gly residues, with an increase in Gly content leading to an increase in linker flexibility.^*40*^ A correlation between linker flexibility and affinity is confirmed by replacement of the entire linker sequence with the highly flexible sequence SGSGS which results in a HIF-1α construct with high binding affinity (I_c/b_ = 0.87), underscoring the need for a dynamic and flexible linker rather than attractive interactions between linker side chains and TAZ1.

### Diverse effects of deletion mutants

AlphaFold models suggest that up to three residues can be deleted from the flexible linker without perturbing the interactions between TAZ1 and the αB and αC regions of the HIF-1α CTAD. A decrease in the linker length might be expected to increase the binding affinity by reducing the entropic cost of binding and enhancing the thermodynamic coupling between the αB and αC motifs.^*37,38*^ However, that is not what is observed: the effect of deletions depends upon which residues, rather than how many, are deleted. Only the deletion of R810, either alone (ΔR) or with S809 (ΔSR) results in an affinity increase, likely due to relief of unfavorable electrostatic interactions with the TAZ1 surface. Single-site deletion of Q807 or S809 has no effect on affinity, while deletion of G808 (ΔG), either singly or with S809 (ΔGS) results in decreased affinity, presumably because of decreased flexibility of the linker and retention of the unfavorable interactions of R810. Shortening of the linker by deletion of three residues can be accommodated, albeit at the cost of reduced binding affinity. Once again the extent of binding impairment depends upon which residues are deleted, with the presence or absence of R810 having a large effect (ΔGSR, I_c/b_ = 0.37; ΔQGS, I_c/b_ = 0.09). Replacement of the WT QGSRN linker with the shorter linker from Coral (*Orbicella faveolata*) impairs binding to a lesser extent (I_c/b_ = 0.19) than the ΔGS mutation (I_c/b_ = 0.05), showing the effect of removing the positive charge by the R810N substitution.

### Effects of linker mutations on the operation of the unidirectional switch

The experiments reported herein show that the affinity of the HIF-1α CTAD for TAZ1 is modulated by changes in the length and composition of the αB - αC linker. Using native gel competition assays, fluorescence anisotropy and NMR measurements, we tested whether these effects translate into changes in the efficiency of the unidirectional switch mediated by CITED2. We have previously proposed a mechanistic model in which the CITED2 αA region interacts transiently with the TAZ1:HIF-1α complex, shifting the conformation of TAZ1 towards that seen in the CITED2-bound state and efficiently displacing the HIF-1α CTAD through competition between the tightly coupled LPEL and hydrophobic ϕ_C_ motifs of CITED2 and the LPQL-αB regions of HIF-1α.^*16,41*^ Rebinding of HIF-1α to the TAZ1:CITED2 complex is strongly inhibited by negative allostery between the CITED2 αA and the HIF-1α αC sites on TAZ1, such that binding at the two sites is almost mutually exclusive.^*17*^ However, the competition between CITED2 and HIF-1α becomes fully reversible when the CITED2 CTD is truncated to remove the ϕ_C_ motif, or when the ϕ_C_ and LPEL motifs are decoupled by substitution of LPEL by a flexible APAA sequence.^*16*^

In addition to the negative allostery associated with conformational changes in TAZ1, we postulated that the 10-residue flexible linker between the HIF-1α αB and αC motifs ensures that the thermodynamic coupling between them is weak, reducing binding cooperativity and further impairing the ability of HIF-1α to compete with and displace TAZ1-bound CITED2.^*10,15*^ The present experiments provide new insights into the role of the linker in determining the directionality of the HIF-1α/CITED2 switch. Mutations and truncations of the HIF-1α linker that increase the binding cooperativity of the αB and αC motifs, as manifest in increased affinity for TAZ1, enable the HIF-1α CTAD to compete more effectively with bound CITED2. Deletion of S809 and R810 (ΔSR), which enhances the binary CTAD binding affinity by 2-fold, results in a variant that is more effective than the WT CTAD at displacing CITED2 from its TAZ1 complex, as observed in both fluorescence anisotropy and NMR competition experiments (Figures 6, 7).

The enhanced competition by ΔSR appears to be associated with loss of R810 rather than decreased linker length since R810G shows similar ability to displace bound CITED2 in native gel assays (Figure S3). Substitution of the central region of the linker with the highly flexible sequence SGSGS also results in a construct with enhanced ability to compete with CITED2. These results underscore the critical role of the conserved R810 in tuning the HIF-1α CTAD binding affinity, decreasing the cooperativity of the αB and αC interactions with TAZ1 to ensure unidirectionality of the switch.

The present work provides new insights into the subtle factors that influence binding avidity of multivalent IDPs. Although the central region (QGSRN) of the αB - αC linker of the HIF-1α CTAD is dynamically disordered and makes no stable contacts with TAZ1, it nevertheless plays an important role in fine-tuning the binding affinity and ensuring unidirectional behavior of the HIF-1α/CITED2 switch. Binding avidity is dominated by the specific intermolecular interactions between the individual binding motifs and the target protein but can be modulated by the length and composition of intervening linkers or intrinsically disordered flanking regions.^*42*^ The conservation of the αB - αC linker length and sequence in all vertebrates suggests strong selective pressure to regulate HIF-1α binding affinity and maintain the unidirectional hypoxic switch; subtle changes in the linker that enhance HIF-1α binding affinity render the switch almost fully reversible. Such selective pressure is absent in organisms that lack CITED2, and the sequence of the αB - αC linker of the linker is highly variable.

## Acknowledgments

We thank Euvel Manlapaz for technical assistance and Gerard Kroon for assistance with NMR experiments. This work was supported by grants GM148226 (PEW) and GM131693 (HJD) from the National Institutes of Health.

## References

[1] Sipko, E. L., Chappell, G. F., and Berlow, R. B. (2024) Multivalency emerges as a common feature of intrinsically disordered protein interactions, Curr. Opin. Struct. Biol. 84, 102742.

[2] Fenton, M., Gregory, E., and Daughdrill, G. (2023) Protein disorder and autoinhibition: The role of multivalency and effective concentration, Curr. Opin. Struct. Biol. 83, 102705.

[3] Weng, J., and Wang, W. (2020) Dynamic multivalent interactions of intrinsically disordered proteins, Curr. Opin. Struct. Biol. 62, 9–13.

[4] Berlow, R. B., Dyson, H. J., and Wright, P. E. (2015) Functional advantages of dynamic protein disorder, FEBS Lett. 589, 2433–2440.

[5] Csizmok, V., Follis, A. V., Kriwacki, R. W., and Forman-Kay, J. D. (2016) Dynamic protein interaction networks and new structural paradigms in signaling, Chem. Rev. 116, 6424–6462.

[6] Jaakkola, P., Mole, D. R., Tian, Y. M., Wilson, M. I., Gielbert, J., Gaskell, S. J., Kriegsheim, A., Hebestreit, H. F., Mukherji, M., Schofield, C. J., Maxwell, P. H., Pugh, C. W., and Ratcliffe, P. J. (2001) Targeting of HIF-a to the von Hippel-Lindau ubiquitylation complex by O_2_-regulated prolyl hydroxylation, Science 292, 468–472.

[7] Arany, Z., Huang, L. E., Eckner, R., Bhattacharya, S., Jiang, C., Goldberg, M. A., Bunn, H. F., and Livingston, D. M. (1996) An essential role for p300/CBP in the cellular response to hypoxia, Proc. Natl. Acad. Sci. U.S.A. 93, 12969–12973.

[8] Kallio, P. J., Okamoto, K., O’Brien, S., Carrero, P., Makino, Y., Tanaka, H., and Poellinger, L. (1998) Signal transduction in hypoxic cells: inducible nuclear translocation and recruitment of the CBP/p300 coactivator by the hypoxia-inducible factor-1a, EMBO J. 17, 6573–6586.

[9] Bhattacharya, S., Michels, C. L., Leung, M. K., Arany, Z. P., Kung, A. L., and Livingston, D. M. (1999) Functional role of p35srj, a novel p300/CBP binding protein, during transactivation by HIF-1, Genes Devel. 13, 64–75.

[10] Berlow, R. B., Dyson, H. J., and Wright, P. E. (2017) Hypersensitive termination of the hypoxic response by a disordered protein switch, Nature 543, 447–451.

[11] Dames, S. A., Martinez-Yamout, M., De Guzman, R. N., Dyson, H. J., and Wright, P. E. (2002) Structural basis for Hif-1a/CBP recognition in the cellular hypoxic response, Proc. Natl. Acad. Sci. USA 99, 5271–5276.

[12] Freedman, S. J., Sun, Z. Y., Poy, F., Kung, A. L., Livingston, D. M., Wagner, G., and Eck, M. J. (2002) Structural basis for recruitment of CBP/p300 by hypoxia-inducible factor-1a, Proc. Natl. Acad. Sci. U.S.A. 99, 5367–5372.

[13] Freedman, S. J., Sun, Z. Y., Kung, A. L., France, D. S., Wagner, G., and Eck, M. J. (2003) Structural basis for negative regulation of hypoxia-inducible factor-1a by CITED2, Nat. Struct. Biol. 10, 504–512.

[14] De Guzman, R. N., Martinez-Yamout, M., Dyson, H. J., and Wright, P. E. (2004) Interaction of the TAZ1 domain of CREB-binding protein with the activation domain of CITED2: regulation by competition between intrinsically unstructured ligands for non-identical binding sites, J. Biol. Chem. 279, 3042–3049.

[15] Berlow, R. B., Martinez-Yamout, M. A., Dyson, H. J., and Wright, P. E. (2019) Role of backbone dynamics in modulating the interactions of disordered ligands with the TAZ1 domain of the CREB-binding protein, Biochemistry 58, 1354–1362.

[16] Berlow, R. B., Dyson, H. J., and Wright, P. E. (2022) Multivalency enables unidirectional switch-like competition between intrinsically disordered proteins, Proc. Natl. Acad. Sci. U.S.A. 119, e2117338119.

[17] Appling, F. D., Berlow, R. B., Stanfield, R. L., Dyson, H. J., and Wright, P. E. (2021) The molecular basis of allostery in a facilitated dissociation process, Structure 29, 1327–1338.

[18] De Guzman, R. N., Wojciak, J. M., Martinez-Yamout, M. A., Dyson, H. J., and Wright, P. E. (2005) CBP/p300 TAZ1 domain forms a structured scaffold for ligand binding, Biochemistry 44, 490–497.

[19] Sugase, K., Landes, M. A., Wright, P. E., and Martinez-Yamout, M. (2008) Overexpression of post-translationally modified peptides in *Escherichia coli* by co-expression with modifying enzymes, Protein Expression and Purification 57, 108–115.

[20] Klock, H. E., and Lesley, S. A. (2009) The Polymerase Incomplete Primer Extension (PIPE) method applied to high-throughput cloning and site-directed mutagenesis, *Meth*. Mol. Biol. 498, 91–103.

[21] Lee, C. W., Ferreon, J. C., Ferreon, A. C., Arai, M., and Wright, P. E. (2010) Graded enhancement of p53 binding to CREB-binding protein (CBP) by multisite phosphorylation, Proc. Natl. Acad. Sci. U.S.A. 107, 19290–19295.

[22] Roehrl, M. H., Wang, J. Y., and Wagner, G. (2004) A general framework for development and data analysis of competitive high-throughput screens for small-molecule inhibitors of protein-protein interactions by fluorescence polarization, Biochemistry 43, 16056–16066.

[23] Delaglio, F., Grzesiek, S., Vuister, G. W., Guang, Z., Pfeifer, J., and Bax, A. (1995) NMRPipe: a multidimensional spectral processing system based on UNIX pipes, J. Biomol. NMR 6, 277–293.

[24] Lee, W., Tonelli, M., and Markley, J. L. (2015) NMRFAM-SPARKY: enhanced software for biomolecular NMR spectroscopy, Bioinformatics 31, 1325–1327.

[25] Hampton-Smith, R. J., Davenport, B. A., Nagarajan, Y., and Peet, D. J. (2019) The conservation and functionality of the oxygen-sensing enzyme Factor Inhibiting HIF (FIH) in non-vertebrates, PLOS ONE 14, e0216134.

[26] Domankevich, V., Opatowsky, Y., Malik, A., Korol, A. B., Frenkel, Z., Manov, I., Avivi, A., and Shams, I. (2016) Adaptive patterns in the p53 protein sequence of the hypoxia- and cancer-tolerant blind mole rat Spalax, BMC Evolutionary Biology 16, 177.

[27] Belato, F. A., Mello, B., Coates, C. J., Halanych, K. M., Brown, F. D., Morandini, A. C., de Moraes Leme, J., Trindade, R. I. F., and Costa-Paiva, E. M. (2024) Divergence time estimates for the hypoxia-inducible factor-1 alpha (HIF1alpha) reveal an ancient emergence of animals in low-oxygen environments, Geobiology 22, e12577.

[28] Mills, D. B., Francis, W. R., Vargas, S., Larsen, M., Elemans, C. P., Canfield, D. E., and Worheide, G. (2018) The last common ancestor of animals lacked the HIF pathway and respired in low-oxygen environments, Elife 7.

[29] Madeira, F., Madhusoodanan, N., Lee, J., Eusebi, A., Niewielska, A., Tivey, A. R. N., Lopez, R., and Butcher, S. (2024) The EMBL-EBI Job Dispatcher sequence analysis tools framework in 2024, Nucleic Acids Res 52, W521–W525.

[30] Abramson, J., Adler, J., Dunger, J., Evans, R., Green, T., Pritzel, A., Ronneberger, O., Willmore, L., Ballard, A. J., Bambrick, J., Bodenstein, S. W., Evans, D. A., Hung, C.-C., O’Neill, M., Reiman, D., Tunyasuvunakool, K., Wu, Z., Žemgulytė, A., Arvaniti, E., Beattie, C., Bertolli, O., Bridgland, A., Cherepanov, A., Congreve, M., Cowen-Rivers, A. I., Cowie, A., Figurnov, M., Fuchs, F. B., Gladman, H., Jain, R., Khan, Y. A., Low, C. M. R., Perlin, K., Potapenko, A., Savy, P., Singh, S., Stecula, A., Thillaisundaram, A., Tong, C., Yakneen, S., Zhong, E. D., Zielinski, M., Žídek, A., Bapst, V., Kohli, P., Jaderberg, M., Hassabis, D., and Jumper, J. M. (2024) Accurate structure prediction of biomolecular interactions with AlphaFold 3, Nature 630, 493–500.

[31] Gu, J., Milligan, J., and Huang, L. E. (2001) Molecular mechanism of hypoxia-inducible factor 1a-p300 interaction. A leucine-rich interface regulated by a single cysteine, J. Biol. Chem. 276, 3550–3554.

[32] Staller, M. V., Ramirez, E., Kotha, S. R., Holehouse, A. S., Pappu, R. V., and Cohen, B. A. (2022) Directed mutational scanning reveals a balance between acidic and hydrophobic residues in strong human activation domains, Cell Syst. 13, 334–345.

[33] Lando, D., Peet, D. J., Gorman, J. J., Whelan, D. A., Whitelaw, M. L., and Bruick, R. K. (2002) FIH-1 is an asparaginyl hydroxylase enzyme that regulates the transcriptional activity of hypoxia-inducible factor, Genes Devel. 16, 1466–1471.

[34] Loenarz, C., Coleman, M. L., Boleininger, A., Schierwater, B., Holland, P. W., Ratcliffe, P. J., and Schofield, C. J. (2011) The hypoxia-inducible transcription factor pathway regulates oxygen sensing in the simplest animal, Trichoplax adhaerens, EMBO Reports 12, 63–70.

[35] Hóbor, F., Hegedüs, Z., Ibarra, A. A., Petrovicz, V. L., Bartlett, G. J., Sessions, R. B., Wilson, A. J., and Edwards, T. A. (2022) Understanding p300-transcription factor interactions using sequence variation and hybridization, RSC Chemical Biology 3, 592–603.

[36] Nyqvist, I., and Dogan, J. (2019) Characterization of the dynamics and the conformational entropy in the binding between TAZ1 and CTAD-HIF-1α, Sci. Rep. 9, 16557.

[37] Krishnamurthy, V. M., Semetey, V., Bracher, P. J., Shen, N., and Whitesides, G. M. (2007) Dependence of effective molarity on linker length for an intramolecular protein-ligand system, J. Am. Chem. Soc. 129, 1312–1320.

[38] Sørensen, C. S., Jendroszek, A., and Kjaergaard, M. (2019) Linker dependence of avidity in multivalent interactions between disordered proteins, J. Mol. Biol. 431, 4784–4795.

[39] Felder, C. E., Prilusky, J., Silman, I., and Sussman, J. L. (2007) A server and database for dipole moments of proteins, Nucl. Acids Res. 35, W512–W521.

[40] van Rosmalen, M., Krom, M., and Merkx, M. (2017) Tuning the flexibility of glycine-serine linkers to allow rational design of multidomain proteins, Biochemistry 56, 6565–6574.

[41] Usui-Ouchi, A., Aguilar, E., Murinello, S., Prins, M., Gantner, M. L., Wright, P. E., Berlow, R. B., and Friedlander, M. (2020) An allosteric peptide inhibitor of HIF-1α regulates hypoxia-induced retinal neovascularization, Proc. Natl. Acad. Sci. U.S.A. 117, 28297–28306.

[42] Karlsson, E., Ottoson, C., Ye, W., Andersson, E., and Jemth, P. (2023) Intrinsically disordered flanking regions increase the affinity of a transcriptional coactivator interaction across vertebrates, Biochemistry 62, 2710–2716.

