## Supplementary material for "Linker Length and Composition within Disordered Binding Motifs modulates the Avidity and Reversibility of a Multivalent Protein Interaction Switch"

**Supplementary Table 1. Sequence Accession Numbers**

| Species | Hif$\alpha$ | CBP | CITED2 |
|---|---|---|---|
| Homo sapiens | NP_001521.1 | NP_004371.2 | NP_001161860.1 |
| Mus musculus | NP_001300848.1 | NP_001020603.1 | NP_001415566.1 |
| Bos taurus | NP_776764.2 | NP_001157494.1 | NP_001069287.1 |
| Equus caballus | XP_023483625.1 | XP_023472189.1 | XP_023488658.1 |
| Canis lupus familiaris | NP_001274092.1 | XP_003434912.1 | XP_038381511.1 |
| Desmodus rotundus | XP_024423159.1 | XP_024409068.1 | XP_024407970.2 |
| Manis pentadactyla | XP_036775412.1 | XP_036776292.2 | XP_036759554.1 |
| Dasypus novemcinctus | XP_058149303.1 | XP_058142543.1 | XP_058163543. |
| Loxodonta africana | XP_010586982.2 | XP_010595656.2 | XP_003404016.2 |
| Phascolarctos cinereus | XP_020842156.1 | XP_020851099. | XP_020829401. |
| Ornithorhynchus anatinus | XP_028935305.1 | XP_028914256.1 | XP_028914380.1 |
| Gallus gallus | NP_989628.2 | XP_015150112.2 | NP_996726.1 |
| Xenopus tropicalis | NP_001011165.2 | NP_001340300.1 | NP_001001196.1 |
| Protopterus annectens | XP_043930427. | XP_043945276.1 | XP_043920657.1 |
| Carcharodon carcharias | XP_041070797.1 | XP_041063371.1 | XP_041064922.1 |
| Mobula hypostoma | XP_062902442.1 | XP_062915522.1 | XP_062911040.1 |
| Latimeria chalumnae | XP_005986474.1 | XP_005990725.1 | XP_006012643.3 |
| Danio rerio | NP_001296971.1 | NP_001352115.1 | NP_001006045.1 |
| Carassius carassius | XP_059368977.1 | XP_059425684.1 | XP_059368401.1 |
| Oncorhynchus mykiss | NP_001117760.1 | XP_036798200.1 | XP_021456139.1 |
| Salmo salar | XP_014066198. | XP_014036978.2 | XP_014060668.1 |
| Petromyzon marinus | XP_032818831.1 | XP_032825942.1 | XP_032828482.1 |
| Branchiostoma floridae | XP_035678467.1 | XP_035682202.1 | XP_035664448.1 |
| Branchiostoma lanceolatum | CAH1263779.1 | XP_066287613.1 | CAH1229276.1 |
| Branchiostoma belcheri | XP_019641152.1 | XP_019625378.1 | XP_019646936.1 |
| Ciona intestinalis | NP_001071731.1; Ensembl ENSCINP00000004277 annotated as Hif1a | XP_018670452. | |

| | | |
|---|---|---|
| Strongylocentrotus purpuratus | XP_030854578.1 | XP_011677839.2 |
| Patella vulgata | XP_050407150.1 | XP_050417373.1 |
| Pecten maximus | XP_033733953.1 | XP_033743941.1 |
| Nematostella vectensis | AII22158.1 | XP_032237806.2 |
| Orbicella faveolata | XP_020630368.1 | XP_020631975.1 |
| Saccoglossus kowalevskii | XP_002733787.2 ; ADB22425.1 | XP_006821556.1 |
| Ptychodera | WNN25270.1 ; XP_070578487.1 | XP_070552633.1 |
| Owenia fusiformis | CAH1791601.1; OQ354998; WNN25263.1 A0AA96K9Z1 | CAH1777193.1 |
| Tribolium castaneum | XP_967427.2 | XP_008192364.2 |
| Procambarus clarkii | XP_069175079.1 | XP_045621111.1 |
| Eriocheir sinensis | UGW01553.1; AHH85804.1 | XP_050718878.1 |
| Rhipicephalus microplus | XP_037274392.1 | XP_037286635.1 |
| Acyrthosiphon pisum | XP_008186097.1 | XP_003242232.1 |
| Parasteatoda tepidariorum | XP_071038973.1 | XP_015922460.1 |
| Apis mellifera | XP_026296721.1 | XP_026294859.1 |
| Diachasma alloeum | XP_015109224.1 | XP_015117846.1 |
| Formica exsecta | XP_029661514.1 | XP_029670996.1 |
| Camponotus floridanus | XP_025263937.1 | XP_011262430.2 |
| Danaus plexippus | XP_061384553.1 | OWR44368.1; XP_061378891.1 |
| Bombyx mori | XP_037876182.1 | XP_062527704.1 |
| Amphimedon queenslandica | XP_011403284.1 no CTAD | XP_019862923.1 |
| Geodia barretti | CAI8029661.1 no CTAD | CAI8019598.1 |
| Trichoplax adhaerens | AFM37575.1; RDD46842.1 no CTAD | RDD39188.1 |

### Supplementary Figures

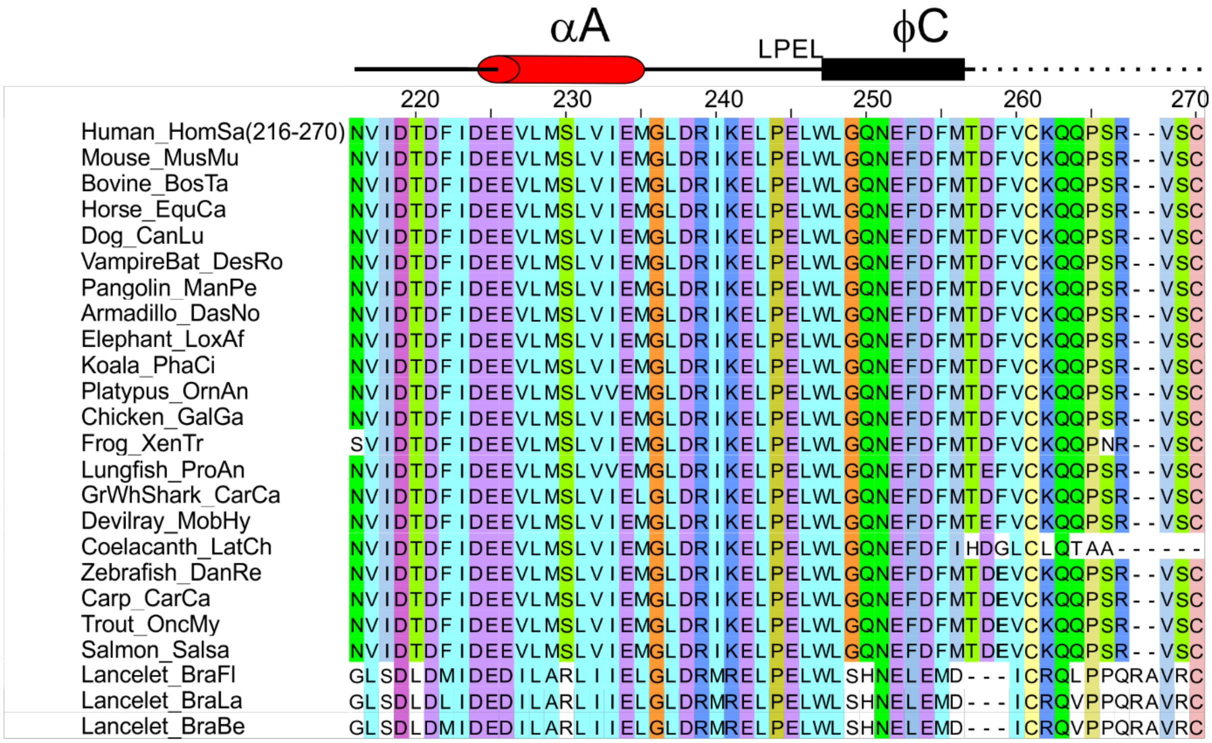

Figure S1a. Alignment of CITED2 TAD sequences from representative chordate species. Binding motifs and secondary structure from the complex with CBP (1R8U) are shown above the alignment.

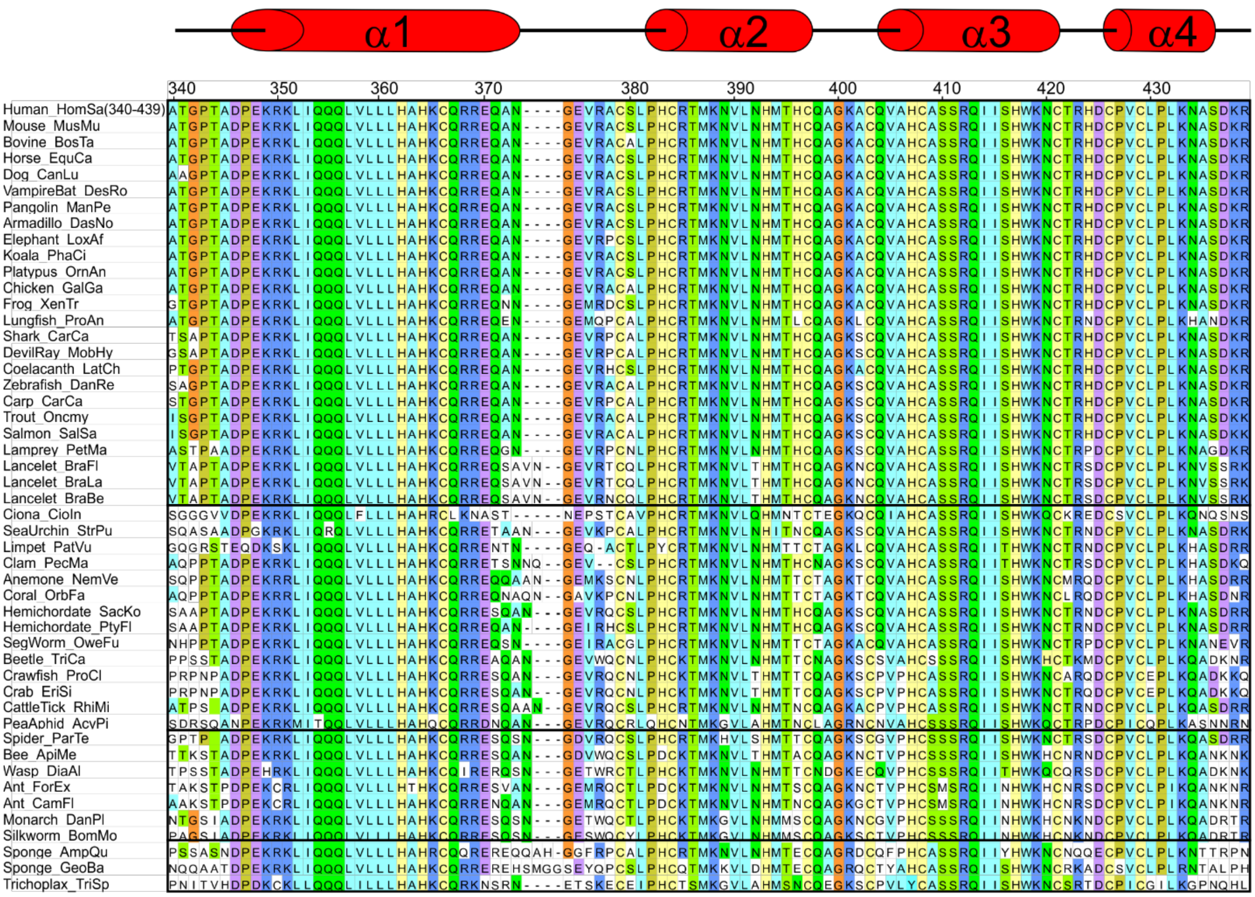

Figure S1b. Alignment of CBP/p300 TAZ1 sequences from representative species. Secondary structure from 1L8C is shown above the alignment. Species in the top group have HIFα, CITED2 and FIH homologues; the second group has HIFα and FIH, the third group has HIFα, and the last group has FIH and a HIFα homologue lacking the CTAD.

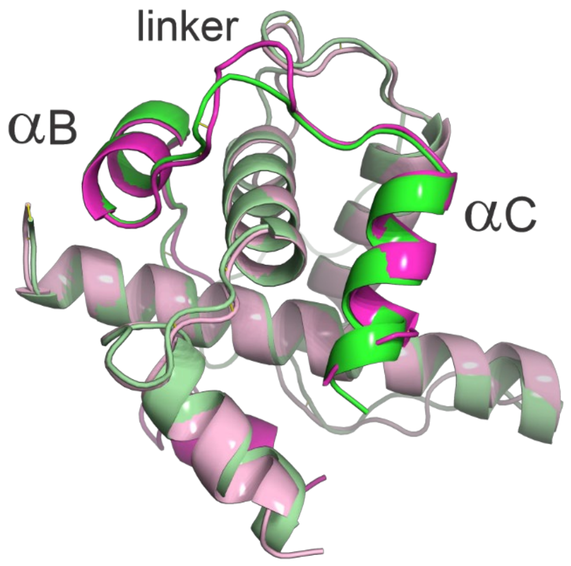

Figure S2. Superposition of AlphaFold3 models of coral_OrbFa HIFα (green) and human HIF-1α (pink) bound to human TAZ1.

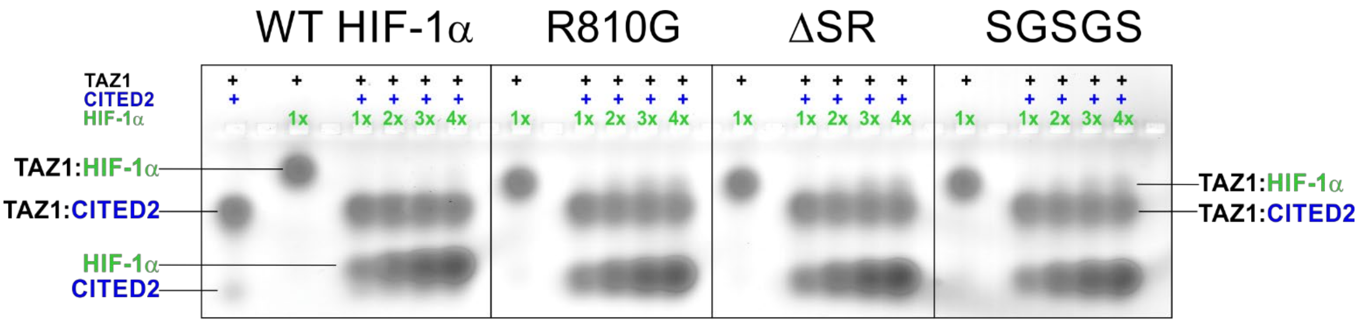

Figure S3. Competition between HIF-1α CTAD high affinity mutants and CITED2 TAD for binding to TAZ1, monitored by native gel electrophoresis. Assay as described in Figure 2 except that in this case both HIF-1α and CITED2 TADs are His6GB1 fusions. TAZ1 and CITED2 TAD are present at 1:1 mole ratio in all samples and HIF-1α CTAD is present at the indicated molar ratio. The mobilities of the HIF-1α and CITED2 TADs in the free and bound states are indicated. A weak TAZ1:HIF-1α band is observable in the lanes containing excess of the HIF-1α variants.

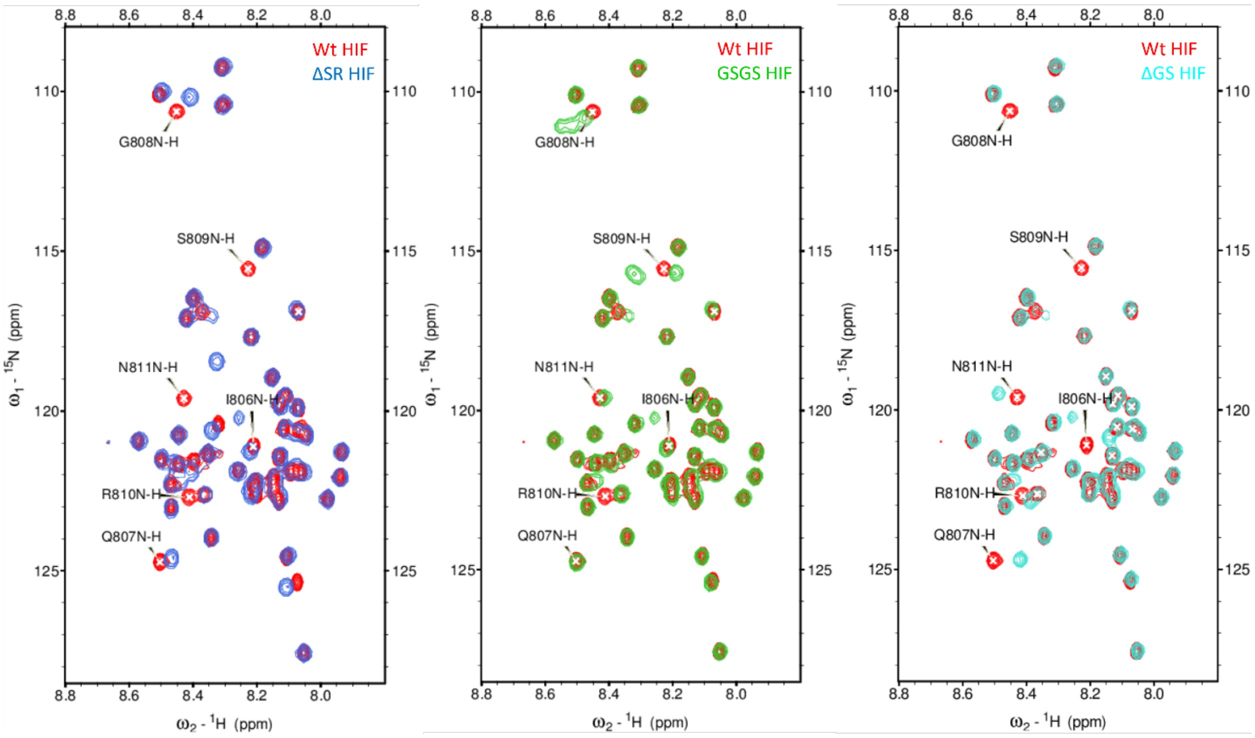

Figure S4: Overlay of $^{15}N$-$^{1}H$ HSQC spectra of free WT HIF-1α CTAD (776-826) (in red) with different HIF variants. ΔSR HIF in blue, GSGS HIF in green and ΔGS HIF in turquoise.

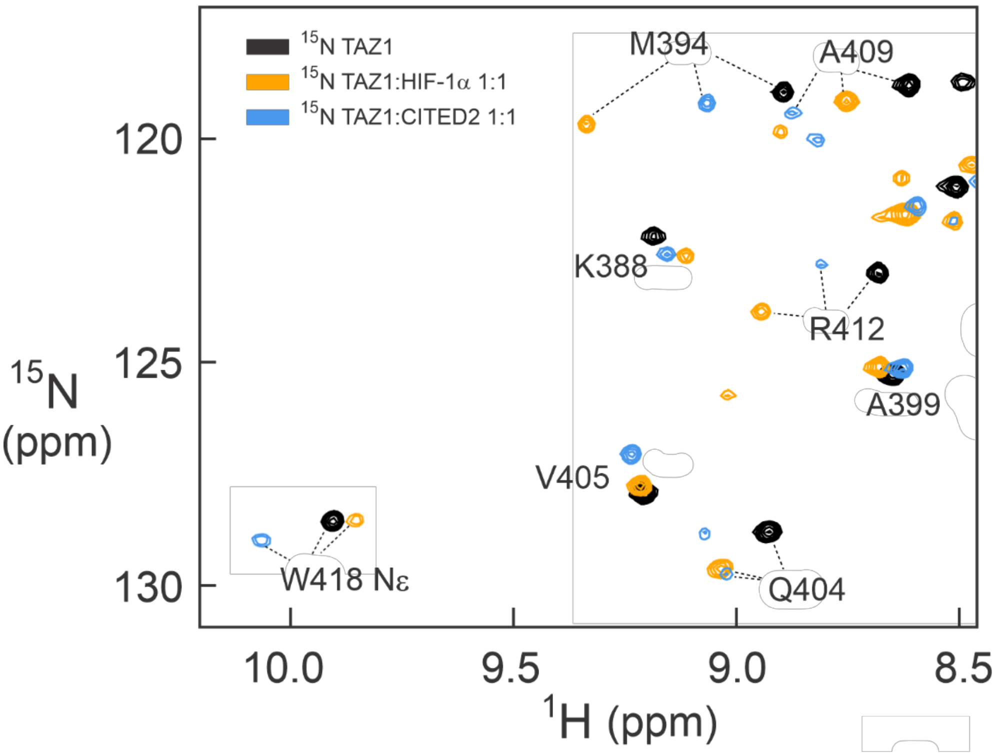

Figure S5. Superposition of portions of the $^{1}H$-$^{15}N$ HSQC spectra of free $^{15}N$-labeled TAZ1 (black), a 1:1 complex of $^{15}N$-labeled TAZ1 with unlabeled HIF-1α CTAD (yellow), and a 1:1 complex of $^{15}N$-labeled TAZ1 with unlabeled CITED2 CTAD (blue). Figure adapted from reference [1] with permission.

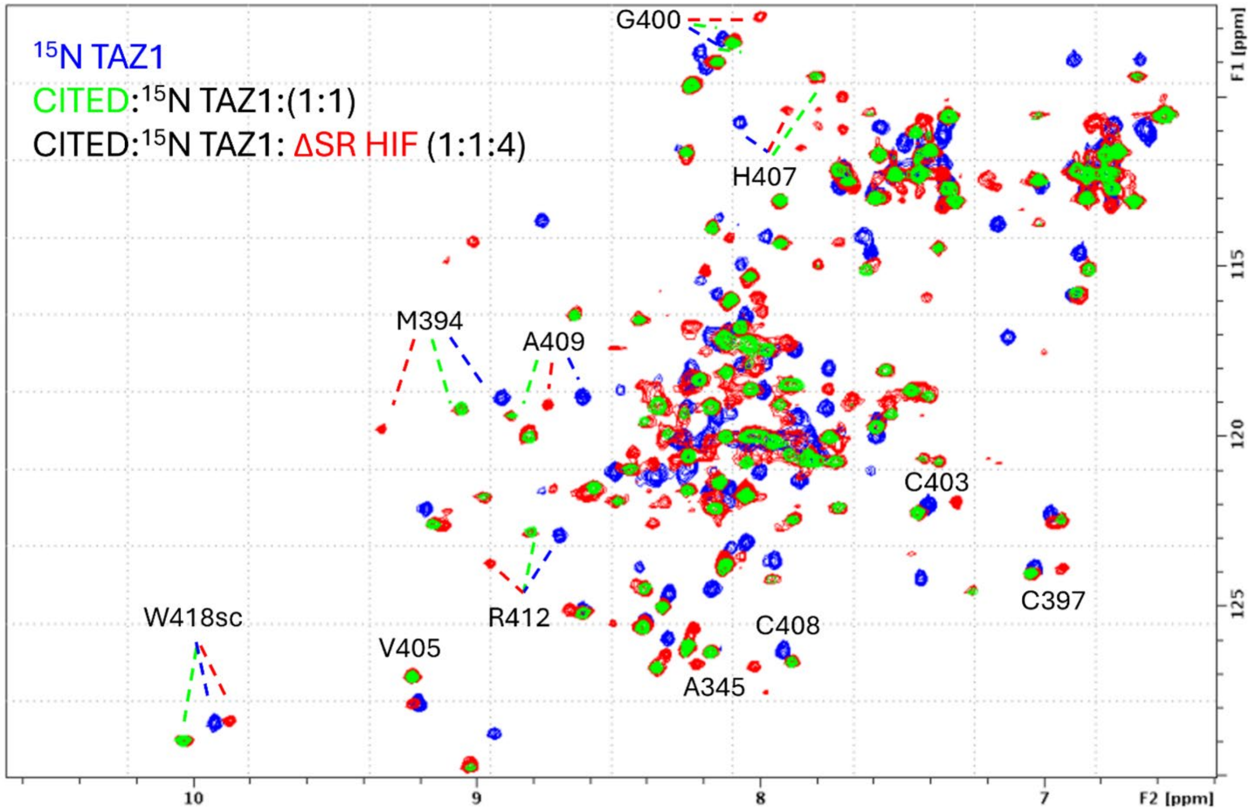

Figure S6: Overlay $^{15}N$-$^{1}H$ HSQC spectra of TAZ1 for free (blue), CITED bound TAZ1 (1:1) (green) and in presence of one molar ratio of CITED and 4 molar equivalents of ΔSR HIF (1:1:4) (red). Selected backbone amide resonances with distinct chemical shifts for the corresponding bound conformations are indicated in similar color arrows.

**Supplementary References**

[1] Berlow, R. B., Dyson, H. J., and Wright, P. E. (2017) Hypersensitive termination of the hypoxic response by a disordered protein switch, *Nature 543*, 447-451.
